# Anticancer Target Combinations: Network-Informed Signaling-Based Approach to Discovery

**DOI:** 10.1101/2024.10.11.617918

**Authors:** Bengi Ruken Yavuz, Hyunbum Jang, Ruth Nussinov

**Affiliations:** Cancer Innovation Laboratory, National Cancer Institute at Frederick, Frederick, MD 21702, USA; Computational Structural Biology Section, Frederick National Laboratory for Cancer Research, Frederick, MD 21702, USA; Department of Human Molecular Genetics and Biochemistry, Sackler School of Medicine, Tel Aviv University, Tel Aviv 69978, Israel

**Keywords:** Combination drugs, combination targets, protein interaction networks, redundant/parallel pathways, compensatory pathways

## Abstract

While anticancer drug discovery has seen dramatic innovations and successes, sequential single therapies are time-limited by resistance, and combinatorial strategies have been lagging. The number of possible drug combinations is vast. To select drug combinations the oncologist requires knowledge of the optimal combination of proteins to co-target. Currently, combinations that the oncologist considers are primarily from empirical observations and clinical praxis. Our aim is to develop a signaling-based method to discover optimal proteins for the oncologist to co-target with drug combinations, and test it on available, patient-derived data. To temper the expected resistance to single drug regimen, we offer a concept-based stratified pipeline aimed at selecting co-targets for drug combinations. Our strategy is unique in its co-target selection being based on signaling pathways. This is significant since in cancer, drug resistance commonly bypasses blocked proteins by wielding alternative, or complementary, routes to execute cell proliferation. Our network-informed signaling-based approach harnesses advanced network concepts and metrics, and our compiled, tissue-specific co-existing mutations. Co-existing driver mutations are common in resistance. Thus, to mimic cancer and counter drug resistance scenarios, our pipeline seeks co-targets that when targeted by drug combinations, can shut off cancer’s *modus operandi*. That is, its parallel or complementary signaling pathways would be blocked. Rotating through combinations could further lessen emerging resistance. We applied it to patient-derived breast and colorectal *ESR1*|*PIK3CA* and *BRAF*|*PIK3CA* subnetworks. Consistently, in breast cancer, our results suggest co-targeting proteins from the *ESR1|PIK3CA* subnetwork with an alpelisib-LJM716 combination. In colorectal cancer, they co-target *BRAF*|*PIK3CA* with alpelisib, cetuximab, and encorafenib combination. Collectively, our pipeline’s results are promising, and validated by patient-based xenografts.

**GRAPHICAL ABSTRACT:** 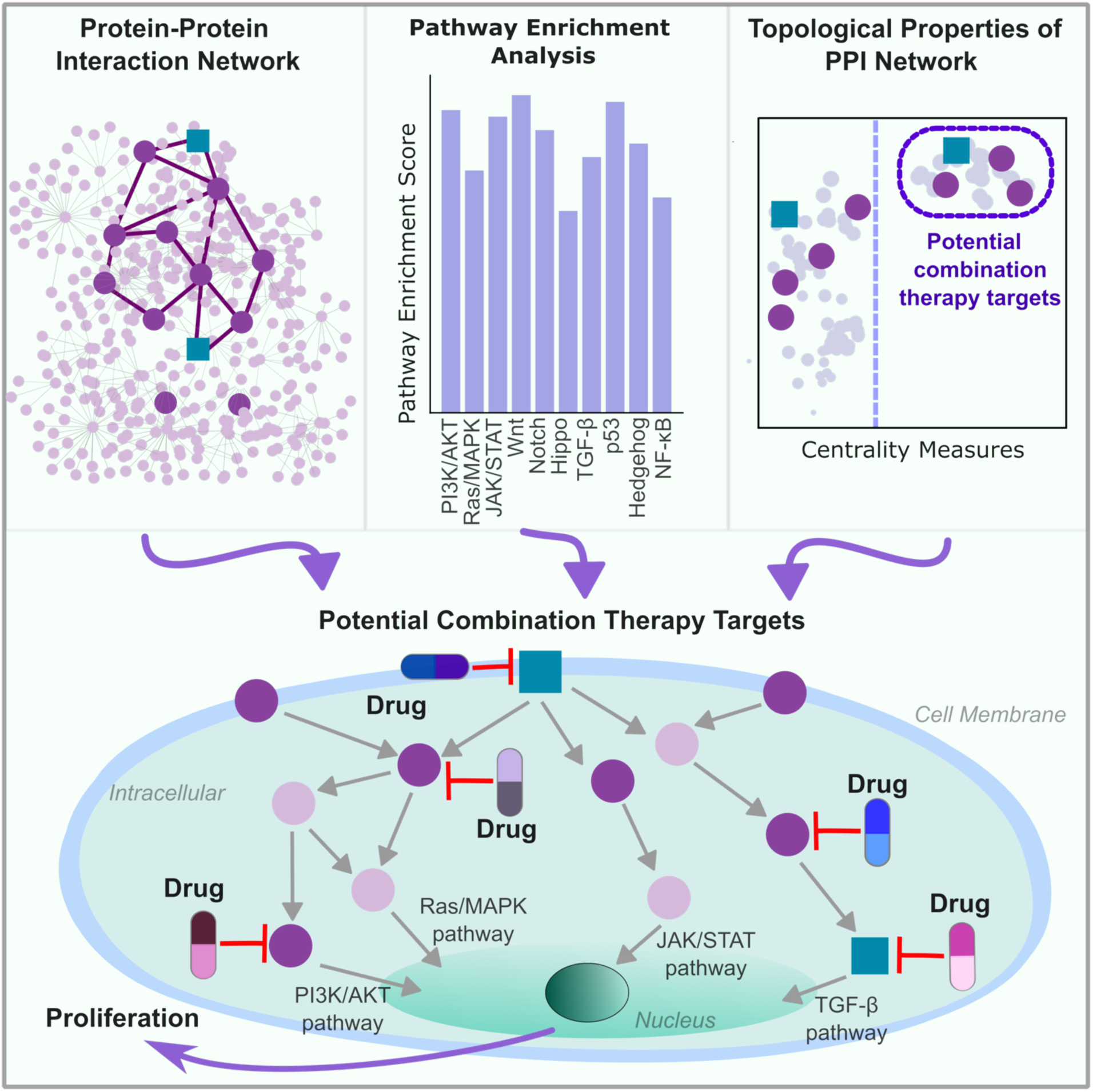

## Introduction

Recent advancements in next-generation DNA sequencing technologies, including single-cell genomics and spatial biology, have dramatically expanded our ability to map complex molecular alterations and epigenetic modifications.^1^ They permit profiling tumor heterogeneity and detection of rare subclonal populations at unprecedented resolution. Integration with multi-omics enable a more holistic understanding of cancer biology, facilitating personalized therapeutics and early detection.^2–7^ However, single drug targeting a single protein (monotherapy) frequently succumbs because of inherent or emergent resistance.^8–10^

Targeting pathways with small molecule inhibitors has been a compelling strategy. Over 80 drugs targeting kinases are FDA-approved, several approved in 2023.^11^ However, cancer cells often develop resistance through alternative growth pathways, with pre-existing drug-resistant subclones proliferating rapidly. One example is PI3K/AKT/mTOR, with around 30-40% of breast cancer cases featuring mutations in *PIK3CA.*^12–17^ The combination of alpelisib, a pan-PI3K inhibitor ^11,18,19^, with hormone therapy has demonstrated effectiveness in metastatic breast cancers that are hormone receptor-positive and human epidermal growth factor (*HER2*) negative.^16,20,21^

Trastuzumab targets patients with HER2-positive breast cancer; however, they may relapse due to activating mutations in *PIK3CA* in 20–30% of the cases. Co-administration of trastuzumab and inhibitors targeting the PI3K/AKT/mTOR signaling axis, including *PIK3CA*, enhances response.^22,23^ In hepatocellular carcinoma, dual inhibition of mTOR and SHP2 shows promising synergistic effects, preventing RTK-mediated resistance to mTOR inhibition.^24^

Pascual et al investigated a phase Ib trial evaluating a novel triplet combination (palbociclib, taselisib, and fulvestrant) in PIK3CA-mutant ER+/HER2-advanced breast cancer and a doublet therapy (palbociclib and taselisib) in PIK3CA-mutant solid tumors.^25^ A notable 37.5% response rate with the triplet therapy in heavily pretreated patients surpassed 15% response rates with palbociclib-fulvestrant alone observed earlier point to the potential of PI3K pathway inhibition.

Methods proposed to optimize combinations include a Graph Convolutional Network (GCN).^26^ Prioritization was also based on semantic relationships between drug and disease^27^, and on pathway crosstalk.^28^ Large scale dual-drug combinations in breast, colorectal, and pancreatic cancer cell lines, revealed the rarity and highly contingent nature of drug synergy.^29^ Combinations of FDA-approved cancer drugs, tested against the NCI-60 cohort,^30^ were deposited at the NCI-ALMANAC repository (A Large Matrix of Anti-Neoplastic Agent Combinations). *In vivo* investigations of murine xenograft models validated the superiority of these drug pairings relative to monotherapies. Finally, NCI-ComboMATCH, augmented the efficacy by genomically directed combinatorial interventions.^31^

Resistance commonly harnesses activating mutations. To identify the mutations, we compiled co-existing, tissue-specific mutations, in the same and in different pathways.^32^ Under the premise that not only the protein pairs harboring co-existing mutations play a critical role in amplifying oncogenic signaling but also proteins on the paths connecting them, we constructed gene-pair specific subnetworks, and identified proteins that serve as bridges between them. The oncogenic subset is comprised of RTKs and TFs, including *EGFR*, *ERBB2, ERBB2*, and *MYC*. These target genes may belong to redundant, parallel, or compensatory pathways.^33^ We validate our computational strategy by application to breast and colorectal cancers. Co-targeting *ESR1*|*PIK3CA* and *BRAF|PIK3CA* were comparable to patient derived xenografts (PDXs), responding better to combination therapies. Co-targeting the *ESR1*|*PIK3CA* subnetwork pathways, a marker of breast cancer metastasis, with alpelisib-LJM716 combination diminished the tumor. Co-targeting *BRAF*|*PIK3CA*, prominent in colorectal cancer, with alpelisib, cetuximab, and encorafenib, which target *PIK3CA*, *EGFR*, and *BRAF* respectively, inhibited tumor growth in colorectal cancer xenografts, suggesting that learning from, and being guided by, nature could be an informed approach to discovery. Here, we refer to protein products of genes (in *italics*), following the source of our data, TCGA and AACR GENIE.

## Results

### Overview of the data and workflow

Precision oncology is increasingly transitioning towards small molecule combinations.^29,33–35^ However, determining the optimal combinations is challenging given the intricate network of molecular interactions and cellular feedback mechanisms.^36–38^ Ideally, all potential drug combinations—double, triple, multiple—should be experimentally evaluated against diverse tumor genomes. However, this is impractical due to their sheer number. Computational methods can significantly narrow the scope of candidates to a manageable scale.

Our primary objective is to delineate the mediators orchestrating intracellular signaling cascades. We define “connector nodes” as communication bridges. In our definition, their interactors are on the shortest cellular network paths. They facilitate efficient signal transduction, essential for rapid cellular responses.^39,40^ Combination therapies targeting them can block oncogenic signals in drug resistance. To engineer the oncogenic signaling landscape, we integrate upstream and downstream regulators in the shortest paths in the subnetworks. For this, we used protein pairs harboring tissue-specific co-existing mutations, taken from our recent pan-cancer genomes study.^32^ We call a protein pair A|B if there is a co-existing mutation pair where the component mutations are on proteins A and B. The shortest paths connect these protein pairs. We construct subnetworks derived from genes on the shortest paths to discern connector nodes. Figure 1 illustrates our pipeline, and the STAR methods section describes the compilation of co-existing mutations,^32^ the computation of the shortest paths in the integrated HIPPIE PPI network^41^ by deploying the Path Linker algorithm,^42^ the pathway enrichment analysis using EnrichR^43^ on a KEGG 2019-derived dataset,^44^ and the gene-pair specific subnetworks using the PageRank algorithm.^45^ We then computationally validate our results using patient-derived xenograft models^46^ for breast cancer and colorectal cancer specific subnetworks. The supplemental materials provide further details of our calculations and the data, including the shortest paths for 1,263 protein pairs harboring the 3424 co-existing mutations (Table S1), and the pathway enrichment analysis (Table S2). Figure S1a-b presents protein pairs where shortest path proteins traverse two and three signaling pathways, respectively. In Figure S1a, protein pairs in the PI3K/AKT pathway combine with either MAPK, Hippo, ErbB, or the Chemokine pathways. The shortest path proteins connecting metastatic markers^32^ are predominantly enriched in PI3K/AKT and MAPK pathways, except for *ESR1/PIK3CA*, enriched in PI3K/AKT and Chemokine pathways. We also show the Ras pathway, deposited as a separate pathway in KEGG data.

**Figure 1.**
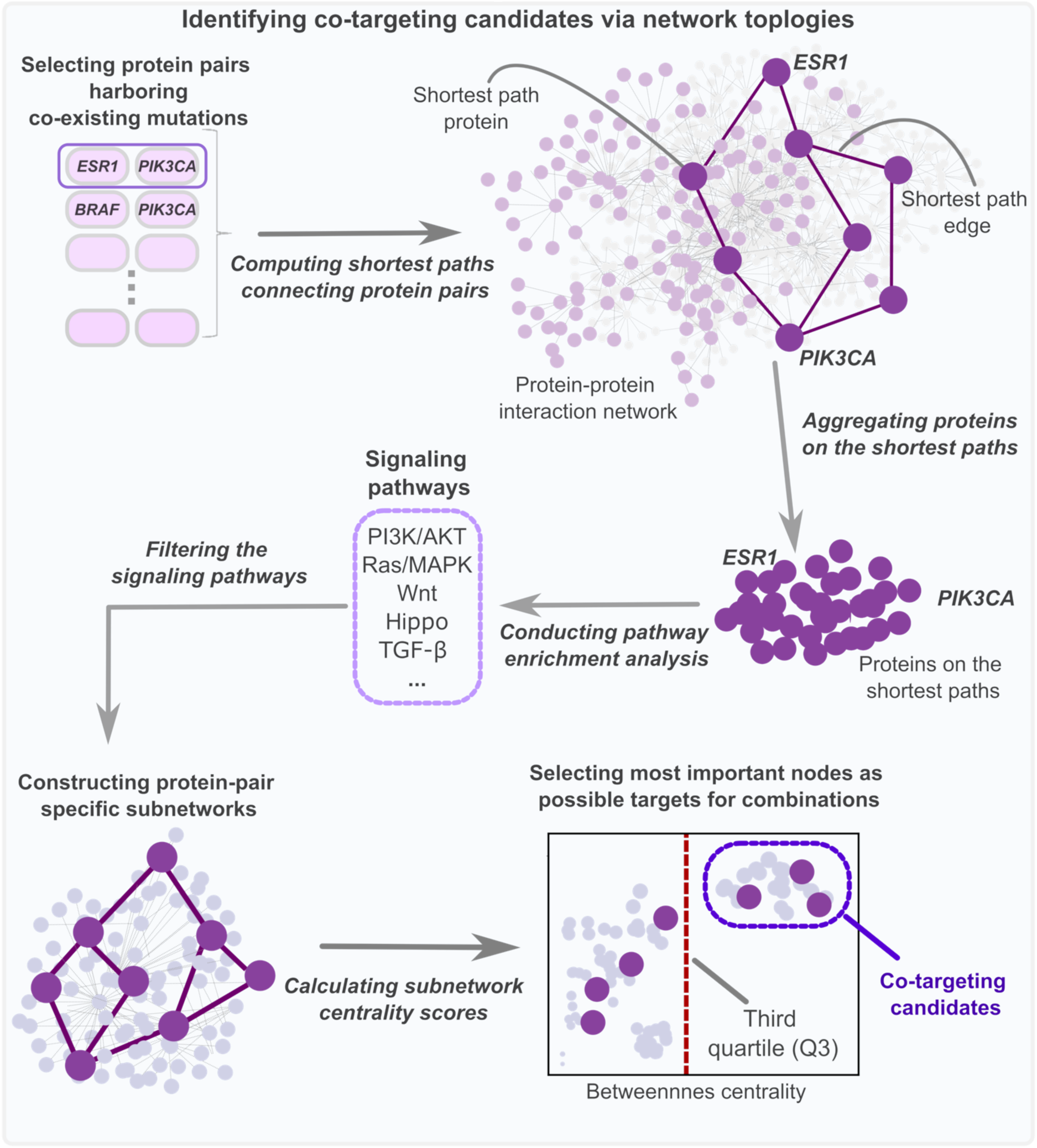
The pipeline of our method aiming to identify critical genes whose encoded proteins harbor co-existing mutations, and whose communication could be important in cancer progression. Our algorithm starts with selecting protein pairs harboring co-existing mutations such as *ESR1*|*PIK3CA*, *BRAF*|*PIK3CA*, etc. Then for each protein pair, we compute the shortest paths connecting them in the HIPPIE ^41^ protein-protein interaction (PPI) network using the Path Linker^42^ algorithm. We aggregate proteins on the shortest paths, and here, conduct pathway enrichment analysis among 46 signaling pathways from KEGG.^44^ We follow with pathway enrichment analysis with EnrichR^43^ focusing on signaling pathways, highlighting those where at least 20% of the shortest path proteins are involved. Using the obtained shortest path proteins in these enriched signaling pathways as seeds for the PageRank^45^ algorithm, we construct protein-pair specific subnetworks. For each such subnetwork, we calculate subnetwork centrality scores identifying those with high betweenness centrality as key connector nodes (bridges) and propose the highest scoring nodes as possible targets for combination therapies, as described below.

We identified 91 protein pairs where more than 20% of shortest path proteins exclusively traverse a single pathway. A bubble plot (Figure S2) visually represents 61 protein pairs, emphasizing those where at least one component is a drug target according to the Connectivity Map (CMAP) data.^47^ Remarkably, a substantial portion of shortest path proteins associated with these pairs navigate through the PI3K/AKT pathway, followed prominently by the MAPK, and thyroid pathways. This underscores the pivotal role of the PI3K/AKT pathway in oncogenic signaling, positioning it as a significant reservoir for identifying multiple drug targets.

For each of the 170 protein pairs, we compose a seed gene set by selecting the shortest path proteins in the enriched pathway(s) to feed into the Page Rank algorithm to reconstruct subnetworks (see STAR Methods). We then calculate and select co-targeting candidate connector nodes (Table S3). Subnetworks constructed from protein pairs in the ‘Single’ pathway category exhibit significantly greater distances compared to ‘Double’ and ‘Multiple’ pathway categories (Figure S3). This suggests that genes in a single pathway are more isolated, connected primarily via a few upstream/downstream proteins, whereas genes in double and multiple pathways exhibit closer proximities, indicating potential crosstalk. In the ensuing sections, we demonstrate the application of our analytical pipeline to the *ESR1|PIK3CA* and *BRAF|PIK3CA* protein pairs, specific to breast and colorectal cancer, respectively.

Deciphering upstream and downstream interactions within pathways, including crosstalks, necessitates discerning their functional similarity in terms of their constituent proteins (Supplemental information). We computed the similarity using the Jaccard index (Figure S4). High pathway proteins similarity suggests the potential for targeting drug interventions upstream and downstream (in the same pathway or in redundant pathways), while dissimilar pathways highlight proteins relating to inter-pathway (parallel or compensatory pathways) crosstalk.^33,48^ For instance, the KEGG MAPK signaling pathway shares significant protein overlap with the PI3K/AKT signaling pathway, indicating high similarity as per the Jaccard Index. Similarly, the ErbB signaling pathway shows substantial overlap with the neurotrophin signaling pathway. In the context of cellular pathways within identical cell types, pathways sharing identical proteins or proteins from analogous families are termed ‘redundant,’ whereas those utilizing distinct proteins are classified as ‘parallel.’ Compensatory pathways are the ones that are in crosstalk.^33^

Understanding how cells communicate is key to identifying the critical drug targets in oncogenic signaling networks. Combination therapies guided by signaling pathways have the potential to inhibit proliferative signaling cascades and detect feedback mechanisms through connector nodes in subnetworks. By examining the pathways through which signals flow and evaluating their similarities, potential drug target combinations can be inferred. We dub important connector nodes “co-targeting candidates”.

### Connector nodes in gene-pair specific subnetworks as co-targeting candidates

We hierarchically clustered a group of genes selected as co-targeting candidates due to their pivotal roles identified by betweenness centrality metrics. Clustering organized these genes based on their signaling pathways. This clustering approach facilitates the selection of targets from each cluster for combination therapies, capitalizing on synergistic interactions inherent to these gene networks. Targeting genes central to essential communication pathways enhances the efficacy of combination therapies compared to single agents.^29,33,35,49^

On the left panel of Figure 2, the dendrogram displays co-targeting candidates. The dendrogram illustrates the relationships, employing the x-axis to depict the Hamming distance metric that quantifies dissimilarity between clusters; smaller distances indicate greater similarity. We constructed a classified dataset with genes as the indices and the signaling pathways the column names. A cell will have value 1 if the corresponding gene is a member of the signaling pathway in the corresponding column, 0 otherwise. Notably, a maximum distance of 0.8 on the dendrogram offers insights into the divergence between clusters. This distance suggests a balance between intra-cluster homogeneity. Greater distances on the dendrogram indicate infrequent co-occurrence of genes within the same signaling pathway, reflecting lower connectivity and functional association. Such separation implies involvement in disparate biological pathways or networks, highlighting divergent cellular roles.

**Figure 2.**
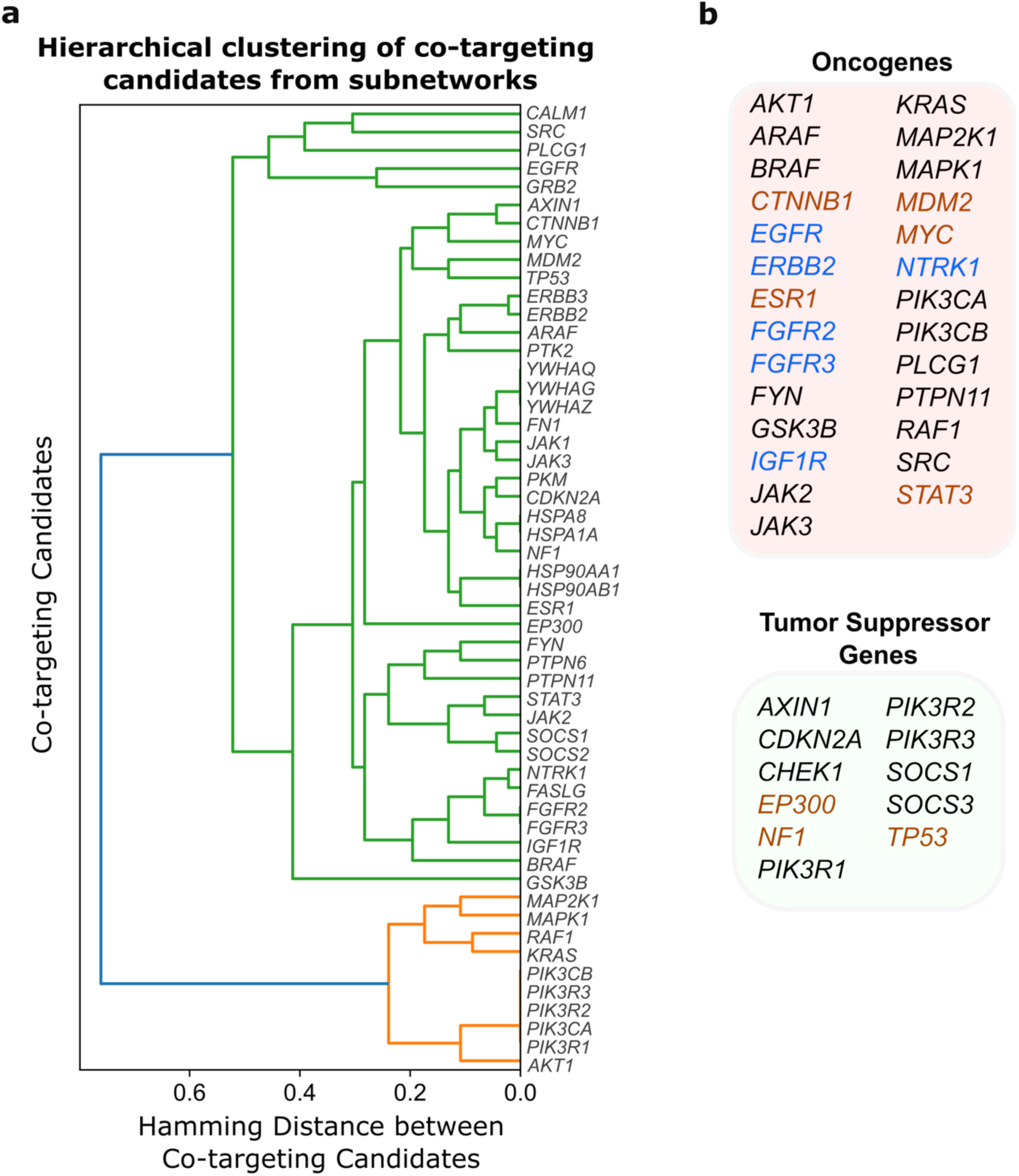
Selection of co-targeting candidate nodes from the subnetworks. **(a)** The dendrogram delineates co-targeting candidate nodes. The y-axis denotes Hamming distance metric that quantifies the dissimilarity between clusters of genes based on their signaling. It measures the number of corresponding nodes in the pathways which are different in the two clusters. Smaller distances indicate higher similarity. Larger distances imply lower similarity. Lower similarity between pathways implies infrequent co-occurrence of genes, suggesting distinct biological pathways or networks in cellular processes. Genes with a Hamming distance of less than 0.1 are deemed functionally similar due to overlapping pathways. *PIK3CA/B* and *PIK3R1/2/3*, with a distance of zero, belong to the same cluster and are in close proximity to *AKT1*. This cluster is connected to the *KRAS* and *RAF* cluster, which is also near the *MAPK1* and *MAP2K1* cluster. Genes positioned farther apart are less likely to share interactions and functions. **(b)** The co-targeting node candidates from (a) are categorized as oncogenes and tumor suppressor genes in boxes on the right. The red and blue fonts are transcription factors (TFs) and the receptor tyrosine kinases (RTKs), respectively. The dataset encompasses 53 co-targeting candidates, including 28 oncogenes and 11 tumor suppressor genes. Among the oncogenes, seven RTKs are identified as co-targeting candidates: *EGFR, ERBB2, ERBB3, FGFR2, FGFR3, IGF1R*, and *NTRK1*. TFs within the oncogenes include *CTNNB1, ESR1, MDM2, MYC*, and *STAT3*.

Genes with Hamming distance <0.1 can be considered as having similar functions as their corresponding pathways overlap. On the right panel of Figure 2, we enlist the oncogenes (OGs) and tumor suppressor genes (TSGs) and the TFs in these groups. The dataset comprises 53 co-targeting candidates composed of 28 oncogenes and 11 tumor suppressor genes. For the oncogenes there are seven RTK co-targeting candidates including *EGFR, ERBB2, ERBB3, FGFR2, FGFR3, IGF1R* and *NTRK1*. As to the TFs among OGs, *CTNNB1*, *ESR1*, *MDM2*, *MYC* and *STAT3* are the co-targeting candidates. We see that *PIK3CA/B, PIK3R1/2/3* (distance 0) belong to the same cluster and this cluster is in proximity to *AKT1*. It connects the clusters of *KRAS* and *RAF* which is also in proximal to the cluster composed of *MAPK1* and *MAP2K1*. Genes positioned farther apart are less likely to share regulatory or interaction networks, emphasizing their distinct functional contexts. Combining proteins from each cluster in therapeutic strategies can enhance treatment efficacy and scope.

Within the PI3K/AKT pathway, several oncogene co-targeting candidates play critical roles. *PIK3CA* and *PIK3CB* encode the p110α and p110β catalytic subunits of PI3K, respectively, which are upstream activators of *AKT1*, a key effector in the PI3K/AKT signaling cascade.^50^ This pathway regulates cellular processes such as growth, metabolism, and survival. Additionally, *GSK3B*, a downstream target of *AKT*, is involved in glycogen metabolism and cell survival, while *CDK4*, regulated by PI3K/AKT signaling, plays a crucial role in cell cycle progression. *PTPN11* (SHP2) also interacts with PI3K signaling, further linking RTK activation to downstream effectors in the PI3K/AKT pathway.^51^ Another pathway that features prominently among the oncogenes identified in the co-targeting set is MAPK. *KRAS, BRAF, ARAF,* and *RAF1* are key upstream regulators in this pathway. *KRAS* acts as an upstream regulator of *RAF* kinases, including *BRAF* and *RAF1*, which subsequently activate MEK1 (*MAP2K1*). MEK1 phosphorylates and activates *ERK* (*MAPK1*), culminating in the regulation of cellular proliferation and differentiation.^52,53^ MAPK is crucial for transmitting signals from the cell surface to the nucleus, transducing gene expression for cell fate decisions. In the JAK/STAT pathway, *JAK2* and *JAK3* encode non-receptor tyrosine kinases involved in cytokine signaling and STAT activation*. STAT3,* a transcription factor activated by JAK kinases, plays a significant role in promoting cell growth and survival, and its constitutive activation is associated with malignancies.^28,54^ Additionally, several oncogenes are involved in other pathways that interface with RTK signaling. *SRC* and *FYN* are members of the Src family of tyrosine kinases, which are involved in various signaling processes, including those related to focal adhesion and RTK signaling.^55^

We next searched the Gene Ontology (GO) biological processes, seeking co-targeting candidates enriched in the Molecular Signatures Database (MSigDB).^56^ The analysis reveals that these co-targeting candidates exhibit significant enrichment in a variety of GO biological processes, notably phosphorylation, apoptosis, and regulation of transferases. These processes are integral to cellular communication and regulatory networks, highlighting the biological significance of the clustered genes. The functional similarities observed across these GO processes suggest that inhibiting these pathways should be considered in therapeutic intervention, as they play pivotal roles in maintaining cellular homeostasis and signal transduction.

Combining RTK inhibitors with inhibitors targeting downstream oncogenes from these pathways aims to achieve a more robust suppression.^57^ One example is co-administering EGFR and PI3K inhibitors, potentially leading to more robust antitumor activity.^58,59^ We propose that genes with a dendrogram distance of less than 0.1 can indicate a shared function of the corresponding genes. Selecting combinations within such clusters can be classified as targeting the same or redundant pathways. Conversely, genes from clusters with a distance of 0.1 or greater can belong to different clusters with distinct functions. Administering combination therapies targeting genes from different clusters can target parallel or compensatory pathways. The critical point here is to map the co-targeting candidates to pathways to identify upstream and downstream proteins, RTKs, oncogenes, and TFs, to infer the signaling mechanisms in a context-dependent manner.

The co-existing mutations network reveals how mutations across proteins contribute to similar phenotypic outcomes through shared biological pathways, highlighting the advantage of considering protein networks over isolated proteins. This method reduces the likelihood of therapeutic resistance, as targeting multiple network nodes diminishes the chance of cancer cell survival. Additionally, addressing interconnected proteins allows combination therapies achieving greater specificity, minimizing off-target effects and preserving cellular functions.

### *ESR1|PIK3CA* subnetwork decodes combination therapy targets for breast cancer

Above, we suggested that the gene sets (encoded proteins) in the Figure 2 dendrogram are important components of cellular communication, and their combinations could be evaluated for combination therapies. To evaluate them as co-targeting candidates, we identified genes in the *ESR1|PIK3CA* specific subnetwork. We reconstructed a subnetwork (Figure 3) specific to metastatic breast cancer by identifying the shortest path proteins that interconnect *ESR1* and *PIK3CA*, which are pivotal markers of breast cancer metastasis.^32^ We discovered that the 200 shortest paths between *ESR1* and *PIK3CA* accommodated 113 proteins in the HIPPIE PPI network and performed pathway enrichment analysis utilizing EnrichR. To delineate the most pertinent pathways associated with the shortest path proteins, we excluded pathways with fewer than 20% of these genes and analyzed the pathway enrichment of the remaining set. For *ESR1*|*PIK3CA*, these pathways are chemokine and PI3K/AKT. For the *ESR1*|*PIK3CA* co-mutated pair, we identified 38 genes that were enriched in the PI3K/AKT and chemokine pathways. 24 proteins were implicated in the chemokine pathway, 30 in the PI3K/AKT signaling pathway, with 16 proteins common to both.

**Figure 3.**
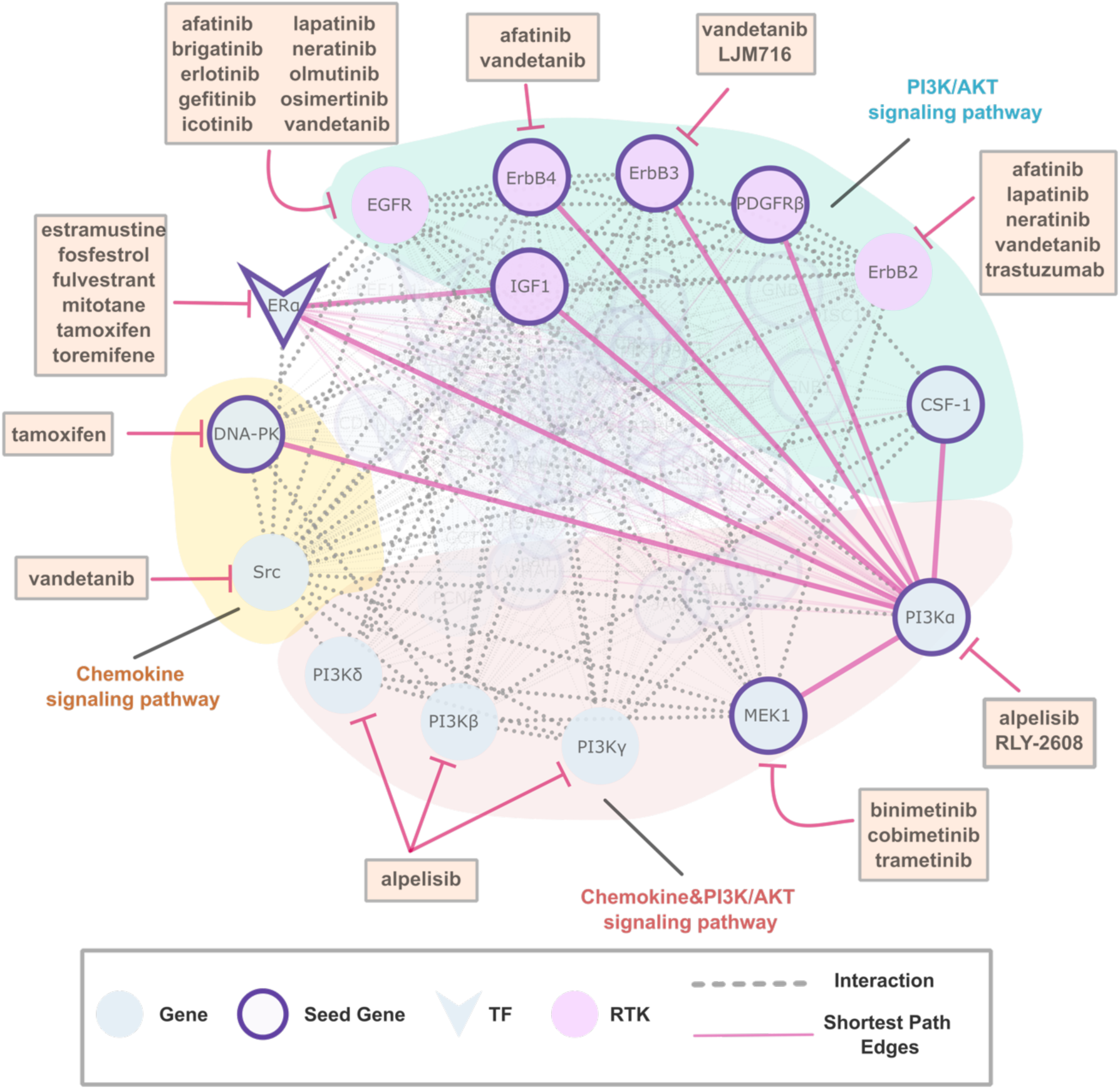
*ESR1|PIK3CA* subnetwork reconstructed with PageRank algorithm in breast cancer. In protein-protein interaction (PPI) networks, a ‘subnetwork’ refers to a smaller, interconnected proteins that work together to regulate specific cellular processes. ‘Subnetwork reconstruction’ refers to the process of identifying and mapping a smaller, functionally relevant group of protein interactions within a larger PPI network. This allows focusing on specific interactions that may play a critical role in cellular functions. This group contributes to the list of co-targeting candidates depending on their betweenness centrality score. We constructed an *ESR1*|*PIK3CA* specific subnetwork using 38 identified shortest path proteins within the chemokine and PI3K/AKT signaling pathways. In protein networks, ‘shortest paths’ refer to the most direct routes (i.e., sequences of interactions) between two proteins within the network. These paths represent the smallest number of interactions (steps) required to connect the two nodes. Key interactions are likely to influence biological processes. Applying the PageRank algorithm, we delineated the *ESR1*|*PIK3CA* specific subnetwork. The resulting subnetwork with 65 nodes provides the drug targets listed in CMAP and RTKs. Key drug targets like *EGFR*, *ERBB2/3/4* (encoding ErbB2/3/4), *PDGFRB* (encoding PDGFRβ), and *IGF1R* (encoding IGF1) are pivotal nodes in the dendrogram, which proposes them as co-targeting candidates. In the reconstructed subnetwork of *ESR1*|*PIK3CA*, seed genes have purple borders, blue nodes are regular genes, and TFs are V-shaped. RTKs are in pink. Inhibitors are in rectangles. The genes in chemokine, PI3K/AKT, and in both pathways are shaded yellow, pink and light blue, respectively.

Then, we reconstructed the *ESR1*|*PIK3CA* specific subnetwork using the 38 identified shortest path proteins as seeds within the chemokine and PI3K/AKT pathways. Utilizing these seed genes, we applied the PageRank algorithm to delineate gene-pair specific subnetworks, excluding proteins with PageRank scores lower than the minimum of either *ESR1* or *PIK3CA*. These subnetworks were designed to elucidate tissue-specific interactions within the HIPPIE PPI network and identify key connector nodes and co-targeting candidates. The resulting subnetwork contains 65 genes with the drug target genes listed in the CMAP. RTKs and shortest path edges are highlighted. The drug target RTKs are *EGFR*, *ERBB2/3/4* (encoding ErbB2/3/4), *PDGFRB* (PDGFRβ), *IGF1R* (IGF1) and the seed gene *CSF1R* (CSF-1) belong to PI3K/AKT pathway. *PIK3CA/B/D/G (*PI3Kα/β/δ/γ) and *MAP2K1* (MEK1) belong to PIK3K/AKT and chemokine pathways, and *SRC* (Src) and *PRKDC* (DNA-PK) belong to chemokine signaling pathway. The transcription factor *ESR1* is not included in these enriched pathways.

We see that the highlighted drug targets in the *ESR1*|*PIK3CA* subnetwork have nodes in the Figure 2 dendrogram, implicating their importance in cellular communication. We also see that *ErbB2* can be targeted with the drugs afatinib, lapatinib, neratinib, vandetanib and trastuzumab. *ErbB3* can be targeted with vandetanib and the monoclonal antibody LJM716, and *EGFR* by several drugs including gefitinib, erlotinib, osimertinib, etc. PI3Kα can be targeted with alpelisib and RLY-2608, an allosteric mutant-selective inhibitor of PI3Kα.^60^ In the subnetwork we also see that *PI3Kβ/δ/γ* are members of both chemokine and PIK3CA/AKT pathways and can be targeted by pan-PI3K inhibitor alpelisib. Another drug target and important mediator is *ESR1* (encoding ERα) which can be targeted by estramustine, fosfestrol, fulvestrant, mitotane, tamoxifen and toremifene from the CMAP data. The clustering in Figure 2 along with the breast cancer specific *ESR1*|*PIK3CA* subnetwork in Figure 3 could be a good strategy for planning combinations targeting one or more RTKs from the oncogenes’ subnetwork (including TFs). In general, if a tumor has activating mutations in at least one of the gene-pair constituents, and seed genes accompanied by sufficient transcriptional activity in the pathways of the seed genes, we can assume an increased oncogenic activity. Either direct inhibition of the connector nodes or indirect inhibition by targeting their partners in protein complexes could provide a venue for combination therapies.

### Co-targeting PI3Kα and ErbB3 in HER2+ breast cancer

In clinical practice, breast cancer is categorized into three therapeutic groups: ER-positive, *HER2* (or ErbB2)-amplified, and triple-negative breast cancers. HER2-positive (HER2+) breast cancer, marked by the overexpression of the human epidermal growth factor receptor 2 (HER2, encoding ErbB2), represents a biologically distinct subset characterized by aggressive tumor behavior and poor prognosis.^61,62^ Targeted therapies, such as trastuzumab and pertuzumab, have markedly improved clinical outcomes for patients with HER2+ breast cancer by specifically inhibiting the HER2 pathway. However, therapeutic resistance remains a significant challenge, often driven by genetic alterations like mutations in the *PIK3CA*.^63^ *PIK3CA* mutations can activate the PI3K/AKT/mTOR pathway, thereby diminishing the efficacy of HER2-targeted therapies by promoting cell survival and proliferation through alternative signaling. Combination therapies that concurrently target HER2 and the PI3K pathways emerged as a promising strategy to overcome resistance.^64^ Combination regimens, incorporating agents such as PI3K inhibitors alongside HER2-targeted drugs, aim to achieve a superior blockade of oncogenic networks, enhancing therapeutic efficacy. The interplay between *HER2* and *PIK3CA* highlights personalized approaches tailored to the molecular profile of the tumor, to optimize outcomes in HER2+ breast cancer.^65–67^

To delineate the signaling mechanism in HER2+ breast cancer, we first map some of the key components in the subnetwork in Figure 3 to the PI3K/AKT pathway (Figure 4). The subnetwork contains the four members of the ErbB family of transmembrane RTKs: EGFR (ErbB1), ErbB2, ErbB3, and ErbB4. ErbB2 and ErbB3 are closely related to EGFR/ErbB1. ErbB2 and ErbB3 frequently form heterodimers, which are highly active in signaling because ErbB2 has potent kinase activity while ErbB3 has an impaired kinase domain but provides strong docking sites for downstream signaling molecules like PI3K.^68,69^ The robust signaling capability of the ligand-activated ErbB2-ErbB3 heterodimer arises from its dual signaling through the Ras/ERK/MAPK pathway, and the PI3K/AKT pathway. This receptor complex evades regulatory downregulation.

**Figure 4.**
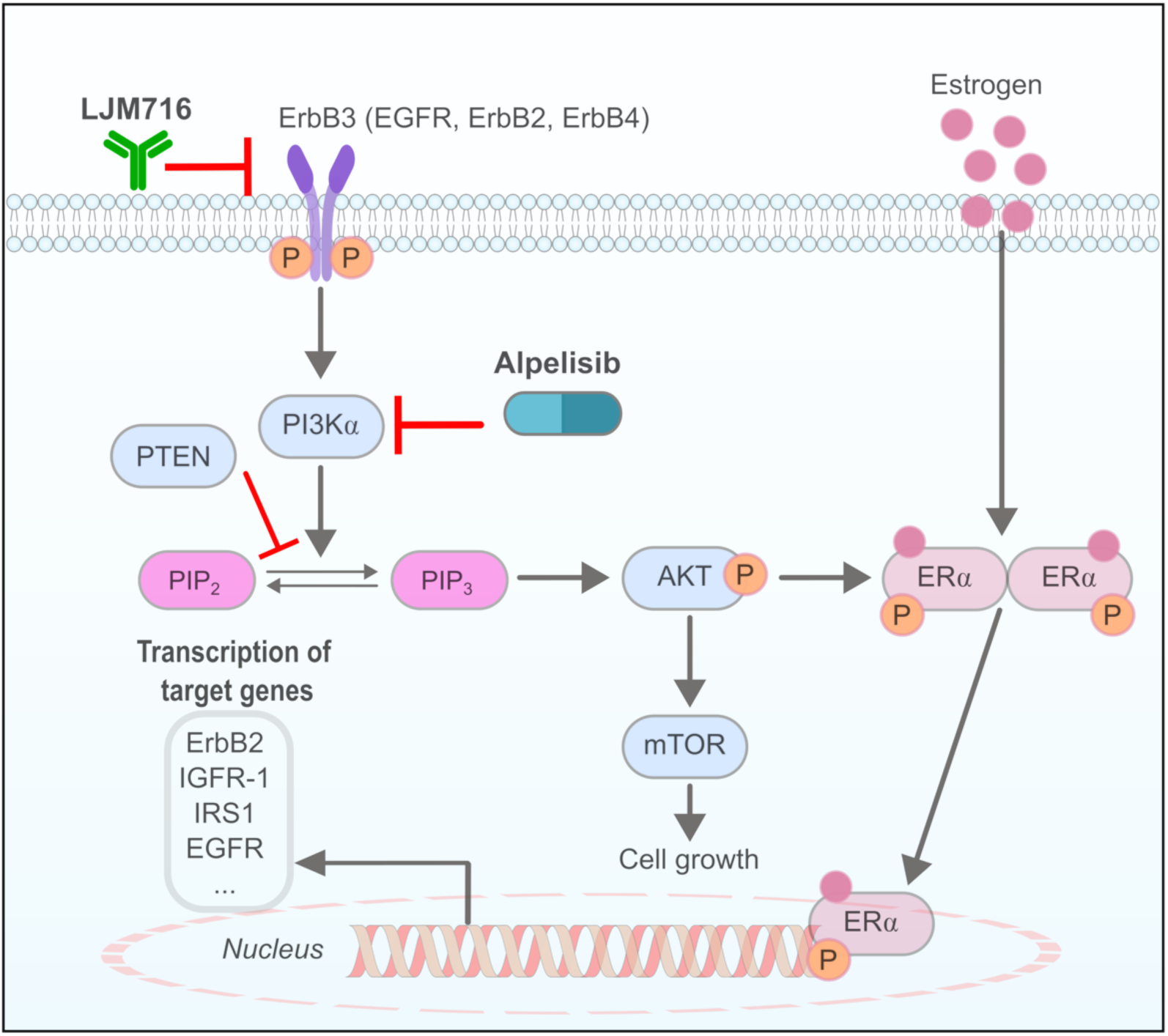
Co-targeting candidates in PI3K/AKT signaling pathway in HER2+ breast cancer. To clarify the signaling mechanism in HER2+ breast cancer, we begin by mapping several key components of the subnetwork onto the PI3K/AKT pathway. The figure is a simple illustration of PI3K/AKT pathway featuring ErbB3 and PI3Kα co-targeted with LJM716 and alpelisib in breast cancer. The combination therapy can block cell growth and transcription of target genes. RTKs such as ErbB3, EGFR, ErbB2 or ErbB4 on the cell surface along with the downstream nodes depict the signaling cascade. Estrogen receptor (ERα) is shown. PI3Kα phosphorylates signaling lipid PIP_2_ to PIP_3_. The ErbB family of RTKs, includes EGFR (ErbB1), ErbB2 (Her2), ErbB3, and ErbB4. ErbB2 and ErbB3 frequently form heterodimers with potent signaling capabilities through the PI3K/AKT pathway.

Upon ligand engagement, the dimer associates with GRB2, either directly through a phospho-tyrosine consensus site or indirectly via SHC interaction, activating MAPK.^68,70,71^ ERK activates transcription factors, including Sp1, Elk-1, and c-Jun. Significant pathways involved are the JAK/STAT and PI3K/AKT. Cyclin-D1, a key regulator of cell cycle progression, is downstream of the ErbB2-ErbB3 heterodimer. Activation of these signaling pathways can induce proliferation, differentiation, migration and apoptosis.^68,69,72^

ErbB2, part of the ErbB receptor tyrosine kinase (RTK) family—comprising EGFR, ErbB3, and ErbB4—operates under stringent spatiotemporal ligand control. Ligand engagement prompts dimerization of ErbB receptors, which activates the cytoplasmic kinase domain. Activation results in tyrosine phosphorylation and the initiation of intracellular signaling pathways.^73,74^ Despite lacking an intrinsic ligand, ErbB2 functions as a coreceptor with ligand-activated ErbBs. ErbB3, characterized by numerous p85/p110 binding sites, excels at activating the PI3K/AKT pathway. However, due to its deficient kinase activity, it necessitates partnering with another ErbB, typically ErbB2, to facilitate the signaling.^73,74^ In breast cancer cells, overexpressed ErbB2 is highly phosphorylated without ligands, causing constitutive activation of the PI3K-AKT pathway. ErbB2 relies on ErbB3 to link to this pathway.^68^ Inhibiting ErbB3 can obstruct formation of ErbB2-ErbB3 heterodimers, attenuating ErbB2-mediated signaling. This inhibition can enhance ErbB2-targeted therapies, such as trastuzumab. Concurrently targeting both partners in the complex, ameliorates inhibition of the pathway. Disruption of ErbB3 affects ErbB2 activity by impeding heterodimerization, reducing signaling through pathways like PI3K/AKT, and likely triggering compensatory mechanisms. This inhibition can be therapeutically advantageous in cancers dependent on ErbB2/ErbB3 signaling.

To identify the nodes pivotal for communication within the subnetwork, we leveraged the networks topological characteristics. We computed the betweenness centrality of proteins within the gene-pair specific subnetwork and determined the third quartile (Q3). We propose genes exhibiting betweenness centrality values exceeding Q3 as key connector nodes (Figure 5a). This analysis revealed several key proteins: *ESR1* (encoding ERα), *PIK3CA* (encoding the protein PI3Kα), *ERBB2* (encoding the protein ErbB2), *IGF1R* (encoding IGF-1), *PIK3R2*, *PIK3R3*, *APP* (amyloid precursor protein), *HCK* (HCK Proto-Oncogene, Src Family Tyrosine Kinase), *FYN* (a membrane-associated tyrosine kinase involved in cell growth regulation and interacting with the p85 subunit of phosphatidylinositol 3-kinase and fyn-binding protein), *EGFR*, and *GRB2 (*which binds to the epidermal growth factor receptor and contains one SH2 domain and two SH3 domains).

**Figure 5.**
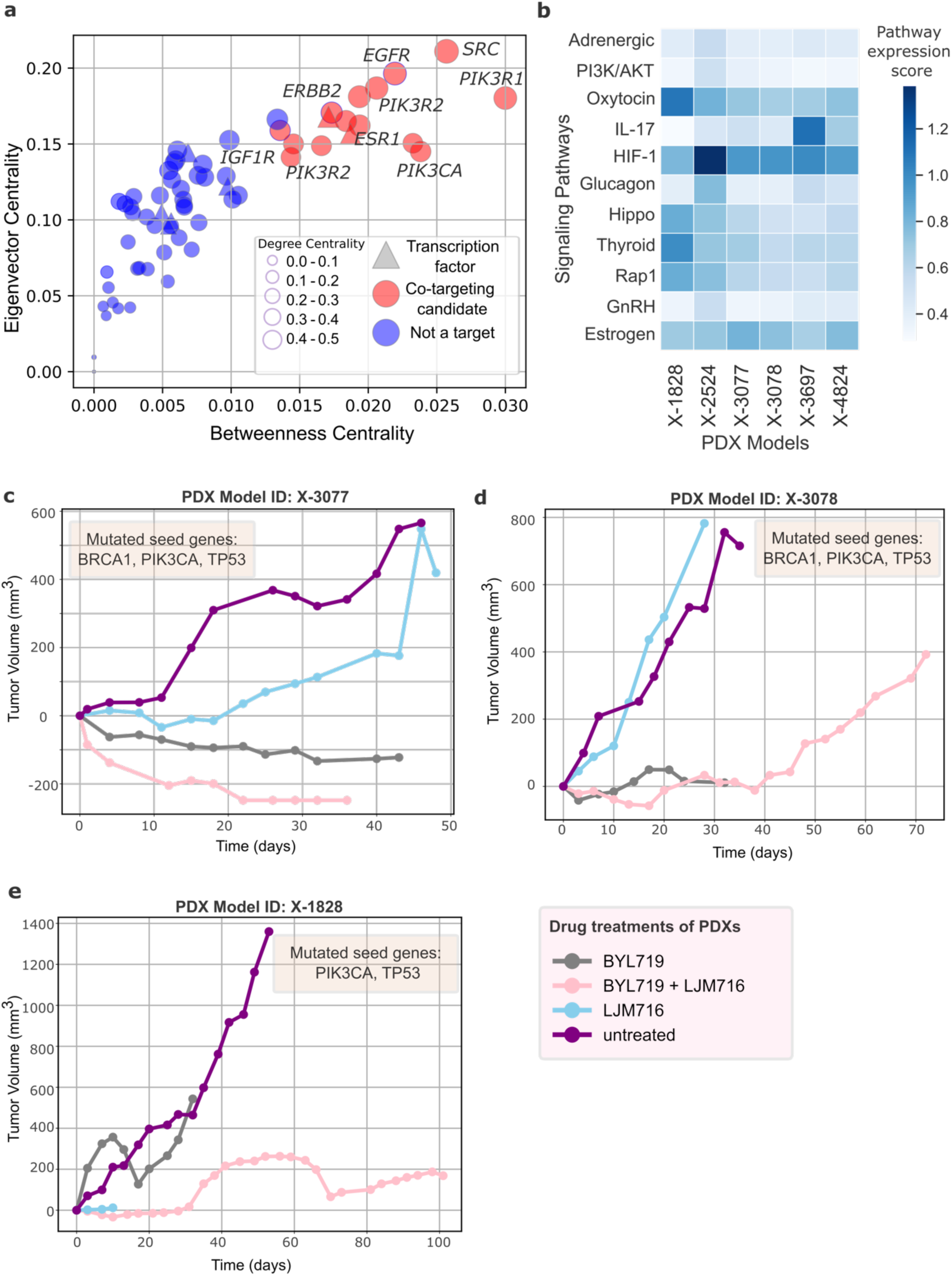
Betweenness centrality and PDX models. **(a)** Scatter plot showing the betweenness centrality (x-axis) and eigenvector centrality (y-axis) of the *ESR1|PIK3CA* subnetwork. Betweenness centrality quantifies the extent to which a node serves as a bridge along the shortest paths between pairs of nodes in a network. Nodes with higher betweenness centrality play crucial roles in efficient communication within the network. Eigenvector centrality measures the influence of a node in a network based on its connections to other central nodes. Nodes with higher eigenvector centrality are not only well-connected but are also connected to other nodes that themselves are highly connected, indicating their influence on the network structure. In the *ESR1*|*PIK3CA* subnetwork, nodes positioned towards the upper right quadrant of the scatter plot exhibit both high betweenness and eigenvector centrality, are co-targeting candidates (red). Nodes that are not potential targets are shown with blue dot. Node sizes are proportional to the degree centrality; nodes with higher degree centrality have more connections, making them potentially more influential in the network. **(b)** Expression scores of upregulated or downregulated pathways in the PDX models with at least one mutation either in *ESR1* or *PIK3CA* and one of the subnetwork genes. There are six such PDX models. All PDXs have high activity in Estrogen, HIF-1 and Oxytocin pathways. **(c-e)** Tumor growth rates of PDX models X-3077, X-3078 and X-1828 when untreated and when treated with BYL719 (alpelisib), LJM716 and BYL719+LJM716. BYL719 is a pan-PI3K inhibitor and LJM716 is anti-HER3 monoclonal antibody. The three PDXs either shrink or grow dramatically slower compared to untreated and monotherapy cases.

Betweenness centrality measures the degree to which a node functions as a bridge in a network. These nodes effectively regulate the flow of communication, as messages typically pass through them. Eigenvector centrality, another significant metric in network analysis, assesses a node’s importance based not only on its direct connections but also on the centrality of its connected nodes. Nodes with high eigenvector centrality are well-connected and linked to other highly central nodes within the network. This metric posits that connections to more central nodes enhance a node’s own centrality score. In terms of communication dynamics, nodes with high eigenvector centrality are influential hubs, extending their impact beyond their immediate vicinity to the entire network.

HER2+ breast cancer can be targeted by combinations of RTKs *EGFR (ERBB1), ERBB2, ERBB3* or *ERBB4* along with a PI3K isoform, *PIK3CA/B/D/G*. In case of *ESR1* mutations or overexpression, *ESR1* can be targeted instead of the RTKs. In Figure 5a (scatter plot of betweenness-eigenvector centrality) we see the important nodes for oncogenic signaling in *ESR1-PIK3CA* network. These include *PIK3CA, ESR1, ERBB2, IGF1R, ERBB2, SRC*, etc. Either targeting these genes directly or interrupting their function through interactor proteins can be a strategy for combination therapies.

For validation, we employed patient-derived xenograft (PDX) models. Gao et al. provided daily drug response rates of PDXs, with volumetric measurements recorded over diverse temporal intervals.^46^ We selected breast cancer (BRCA) models featuring at least two mutations: one in the protein pair components and the other(s) in the seed genes (shortest path proteins). Additionally, to facilitate comparative analysis, we chose models that had temporal volume data for untreated tumors and those treated with combination therapies as well as the individual components of these combinations. This selection resulted in six BRCA models with volume changes recorded over various time intervals for the untreated cases, those treated with BYL719 (alpelisib, targeting *PIK3CA*), LJM716 (an anti-HER3 antibody), and their combination BYL719+LJM716. We provide the expression values BRCA PDX seed genes and drug targets expression values to determine the breast cancer subtype of the models. As both ErbB2 (HER2) and ErbB3 (HER3) levels are relatively higher, we can evaluate these models of HER2+ type (Figure S5). In Figure 6b, we present pathway expression scores for pathways with relatively high expression (Methods). This reveals which pathways exhibit higher expression across all models; notably, the expression scores for the Oxytocin, HIF-1, and Estrogen signaling pathways are elevated. Furthermore, for models X-1828 and X-2524, the scores for Hippo, Thyroid, and Rap1 pathways are also elevated. Regarding the mutation profiles of these models, X-3077 and X-3078 harbor *BRCA, PIK3CA*, and *TP53* mutations among the seed genes, while X-1828 contains *TP53* and *PIK3CA* mutations.

**Figure 6.**
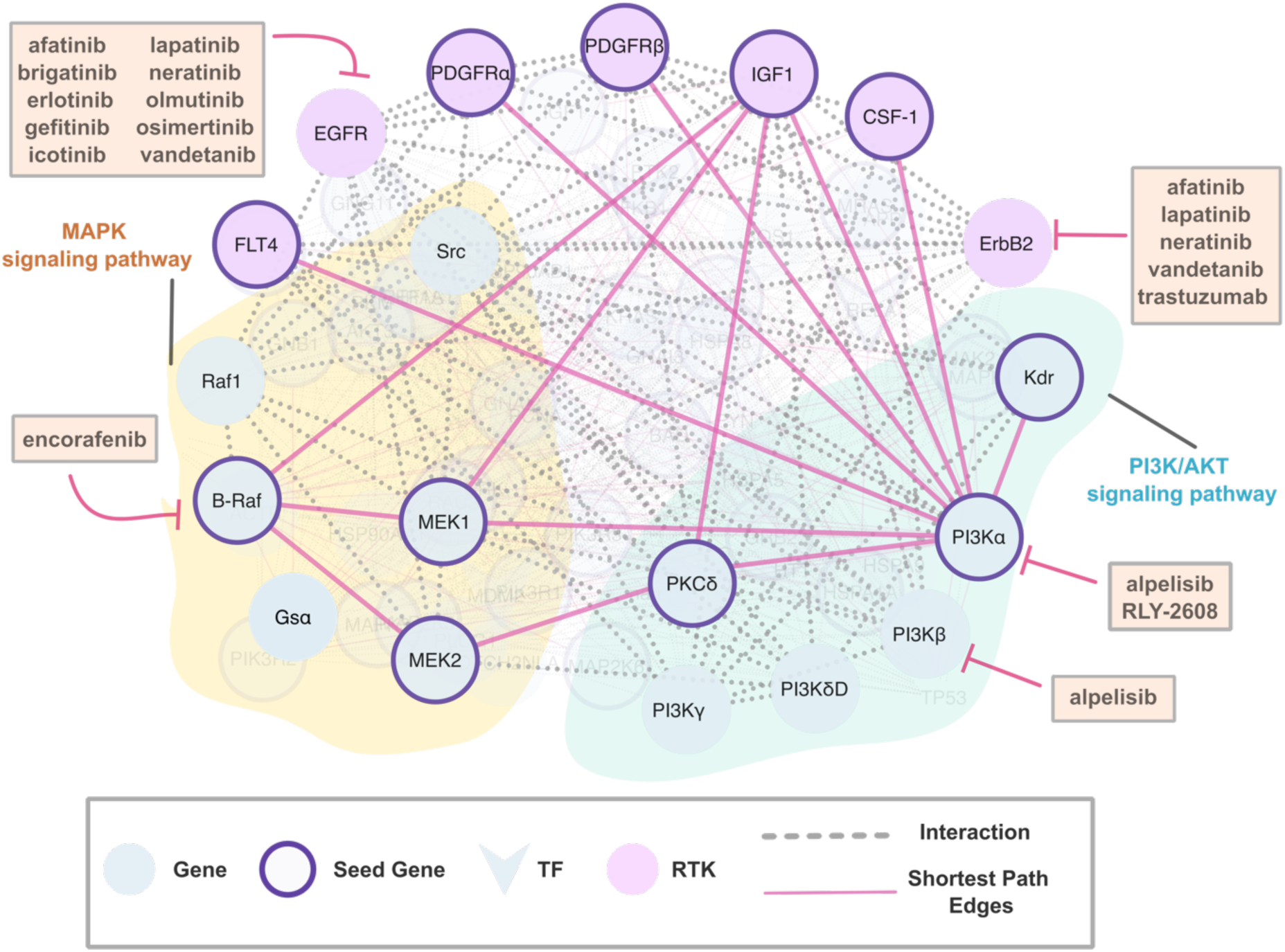
*BRAF|PIK3CA* subnetwork in colorectal cancer reconstructed with PageRank algorithm. *BRAF*|*PIK3CA* subnetwork nodes in the MAPK (in yellow) and PI3K/AKT (blue) pathways delineate a functionally relevant cluster of protein interactions derived from the shortest path proteins linking *BRAF* and *PIK3CA*. Shortest paths refer to the most efficient routes (sequences) of interactions between two nodes within the network. These paths represent the minimal number of interaction steps required to connect the two nodes. The subnetwork of proteins on the shortest paths encompasses RTKs such as *EGFR*, PDGFRα, IGF1, and ErbB2, alongside downstream B-Raf and *PI3K*α, within the compensatory MAPK and PI3K/AKT cascades. The network suggests that co-targeting these RTKs, in conjunction with at least one of the identified oncogenes, offers robust therapeutic strategies as they are the key bridging nodes. These key nodes with high betweenness centrality listed among the co-targeting candidates. In this subnetwork, seed genes are marked with purple borders, regular genes as blue nodes, and TFs are V-shaped. RTKs are in pink. Inhibitors targeting specific nodes within this subnetwork are annotated within rectangular labels.

The growth patterns of models X-3077, X-3078, and X-1828 are illustrated in Figure 5c-e. All three graphs indicate that untreated tumor growth is highly aggressive. In model X-3077, treatment with LJM716 decelerates tumor growth; however, resistance emerges after day 0. Similarly, in model X-3078, resistance to LJM716 becomes evident after day 10. For both X-3077 and X-3078, BYL719 treatment nearly halts tumor growth. Conversely, in model X-1828, the tumor continues to grow despite an initial shrinkage with BYL719 treatment. Under the combination therapy BYL719+LJM716, X-3077 experiences cessation and subsequent shrinkage of tumor growth. Model X-3078 initially responds well to the combination therapy for the first 40 days but begins to grow afterward, though the growth rate remains lower compared to the untreated case. Model X-1828 shows responsiveness to the combination therapy during the first 30 days and after 70 days, with a minor tumor growth observed between days 30 and 70. We also show another model’s response to the drugs in Figure S6. Similarly, PDX models with the metastatic marker mutations of *CDH1* and *PIK3CA* respond better to BYL719+LJM716 and tumor growth slows down dramatically (Figure S7-S8).

Combination therapies work better: Our pursuit of combination therapies is bolstered by the growth rates of patient-derived xenografts (PDXs) under both untreated conditions and various monotherapy and combination therapy regimen (Figure S9). We selectively included 223 models with documented volume data from day 0 and observable growth trajectories in untreated scenarios. We conducted a rigorous analysis of the daily growth rates for these models when administered 24 combination therapies and 17 monotherapies, which are constituent elements of the combination regimens.

Breast cancer cells with overexpressed ErbB2 rely on its activity for proliferation, as treatment with ErbB2-specific antagonistic antibodies or kinase inhibitors arrests these tumor cells in the G1 phase of the cell cycle. Notably, the cessation of ErbB2 signaling correlates with reduction in the phosphotyrosine levels of ErbB3. These findings suggest that ErbB3 may serve as a collaborative partner with ErbB2 in facilitating cellular transformation. Overexpression and activity of ErbB2 alone are insufficient to drive breast tumor cell division. The oncogenic RTK *ERBB2* is recognized for its tendency to dimerize with other members of the EGFR family, particularly ERBB3, robustly activating PI3K signaling. Over two decades, antibody-mediated inhibition targeting the ErbB2/ErbB3/PI3K axis has been a primary treatment approach for patients with *ErbB2*-amplified breast cancer. However, lack of therapeutic response and rapid onset of relapse raised concerns about the assumption that the *ERBB2/ERBB3* heterodimer is the sole relevant effector target.^75^ Inhibiting *ErbB3* signaling can induce non-canonical ErbB2 heterodimers, promoting ErbB2-mediated activation of the ERK pathway.

We evaluated the efficacy of LJM716, a HER3-neutralizing antibody, as monotherapy and in conjunction with BYL719, a p110α-specific inhibitor, in HER2-overexpressing breast and gastric carcinomas. LJM716 effectively diminished HER2-HER3 and HER3-p85 dimerization and phosphorylation of *HER3* and *AKT*, inhibiting tumor progression in xenograft models. The combination of LJM716 with lapatinib and trastuzumab significantly enhanced survival outcomes.

Synergistic interaction between LJM716 and BYL719 resulted in greater inhibition of cell proliferation and AKT phosphorylation in HER2+ and PIK3CA mutant cell lines, surpassing monotherapies. In trastuzumab-resistant HER2+/PIK3CA mutant MDA453 xenografts, combination therapy achieved complete tumor regression, in contrast to partial inhibition observed with individual agents. Long-term treatment showed that only 14% of mice receiving the LJM716/BYL719/trastuzumab combination experienced tumor recurrence, a significantly lower rate compared to other treatment groups. These findings suggest that dual inhibition of HER2 signaling with LJM716 and BYL719, absent direct HER2 antagonism, represents an effective approach for HER2-overexpressing malignancies.^76^

Targeted therapies have markedly transformed the treatment paradigm for estrogen receptor-positive (ER+) breast cancer by precisely targeting molecular pathways crucial for tumor proliferation and survival. Central to this paradigm are mutations in ESR1 and PIK3CA, which frequently co-occur and significantly influence transcriptional processes that regulate cellular proliferation and survival pathways. *ESR1* mutations can induce estrogen receptor activity independent of estrogen. PI3Kα mutations activate the PI3K/AKT/mTOR pathway, thereby enhancing cellular growth, metabolic functions, and survival mechanisms. The interplay between *ESR1* and *PIK3CA* mutations underscores their synergistic impact on transcriptional regulation, amplifying oncogenic signaling pathways in ER+ breast cancer. Understanding the complex molecular mechanisms underlying these mutations is crucial for developing targeted therapies.

Therefore, HER2+ breast cancer can be effectively targeted using combinations of RTKs such as *EGFR (ERBB1), ERBB2, ERBB3,* or *ERBB4* in conjunction with one of the PI3K isoforms, including *PIK3CA, PIK3CB, PIK3CD* or *PIK3CG*. In instances of *ESR1* mutations or overexpression, targeting *ESR1* itself or in combination with other RTKs and with a downstream protein could be a good strategy.

### Combination therapy for colorectal cancer through *BRAF*|*PIK3CA* specific subnetwork

Colorectal cancer (CRC) represents a heterogeneous malignancy characterized by diverse genetic alterations that drive tumorigenesis and progression. Among these aberrations, co-existing mutations in *BRAF* (encoding B-Raf) and *PIK3CA* have garnered significant attention due to their implications for prognosis and therapeutic response. B-Raf, a component of the MAPK signaling pathway, and PI3Kα, a pivotal regulator of the PI3K/AKT pathway, play critical roles in cell proliferation, survival, and differentiation. Biomarkers guided colorectal cancer treatment. Knowledge of primary resistance mechanisms could facilitate potent drug combinations, particularly mutations in the MAPK pathway pertinent to EGFR-targeted and/or B-Raf-targeted treatments.^77^ We aim to identify key nodes in the communication network by setting our starting point at B-Raf and PI3Kα, two genes harboring co-existing mutations specific to CRC.^32^

To this end, we identified genes in the *BRAF|PIK3CA* specific subnetwork. We reconstructed a subnetwork specific to metastatic breast cancer by identifying the shortest path proteins that connect *BRAF* and *PIK3CA*, which are pivotal markers of breast cancer metastasis.^32^ From an analysis of the 200 shortest paths between nodes in the HIPPIE PPI network, we identified 150 proteins involved in these pathways. Using EnrichR for pathway enrichment analysis, we focused on 46 curated KEGG signaling pathways. Pathway enrichment analysis revealed key signaling pathways with over 20% of the shortest path proteins, including MAPK, PI3K/AKT, and Chemokine signaling for *BRAF*|*PIK3CA*. In total, 35 genes were enriched in these pathways for the *BRAF*|*PIK3CA* co-mutated pair.

We constructed *BRAF*|*PIK3CA* specific subnetwork using the 38 shortest path proteins as seeds within the MAPK and PI3K/AKT signaling pathways. Using these seed genes, we applied the PageRank algorithm to delineate gene-pair-specific subnetworks, excluding proteins with PageRank scores lower than the minimum of either *BRAF* or *PIK3CA*. These subnetworks were intended to elucidate tissue-specific interactions within the HIPPIE PPI network and to identify key connector nodes and co-targeting candidates. The resulting subnetwork contains 65 genes (Figure 6) with the drug target genes *listed in the* CMAP, RTKs, shortest path edges are highlighted. In the subnetwork, the drug target RTKs are *CSF1R* (encoding CSF-1), *EGFR* (EGFR), *ERBB2* (ErbB2), FLT4 (FLT4), *IGF1R* (IGF1), *PDGFRA/B* (PDGFRα/β). Downstream in these pathways, we see MAPK pathway members *BRAF* (B-Raf)*, RAF1* (Raf1), *GNAS* (GSα) and *MAP2K1/2* (MEK1and MEK2). From the PI3K/AKT pathway we see PI3K isoforms *PIK3CA/B/D/G* (PI3Kα/β/δ/γ) and two other genes *PRKDC* (encoding PKCδ) and *KDR* (Kdr).

We see that the highlighted drug targets in the *BRAF|PIK3CA* subnetwork have nodes in the dendrogram in Figure 2 implicating their importance in cellular communication. We see that the RTK *ErbB2* can be targeted with afatinib, lapatinib, neratinib, vandetanib and trastuzumab. *EGFR* can be targeted by several drugs including cetuximab, gefitinib, erlotinib, osimertinib, etc. B-Raf can be targeted by encorafenib and PI3Kα/β/δ/γ can be targeted by pan-PI3K inhibitor alpelisib. The principal co-targeting candidates depicted in Figure 2, derived from co-existing mutations with shortest path proteins enriched in at least three pathways, in conjunction with the *BRAF*|*PIK3CA*-specific subnetwork illustrated in Figure 6, can facilitate the design of combination therapies. In general, if a tumor has activating mutations in at least one of the gene-pair constituents and seed genes accompanied with sufficient transcriptional activity in the corresponding pathways of the seed genes, we may assume an increased oncogenic activity mediated by the reconstructed subnetwork nodes. Thus, either direct inhibition of the connector nodes or indirect inhibition by targeting their partners in protein complexes could provide a venue for combination therapies.

The communication network between *BRAF* and *PIK3CA* spans five KEGG signaling pathways: MAPK, Ras, Rap1, PI3K/AKT, and chemokine. Ras and MAPK are upstream and downstream the same signaling cascade and their protein members are similar as shown by Jaccard index similarity (Figure S4). We then merge the two pathways and design a combination therapy by targeting one of the RTKs from the subnetwork and two members from the MAPK and PI3K/AKT pathways (Figure 7a). This strategy can aid in identifying novel combination therapies for targeting parallel pathways.

**Figure 7.**
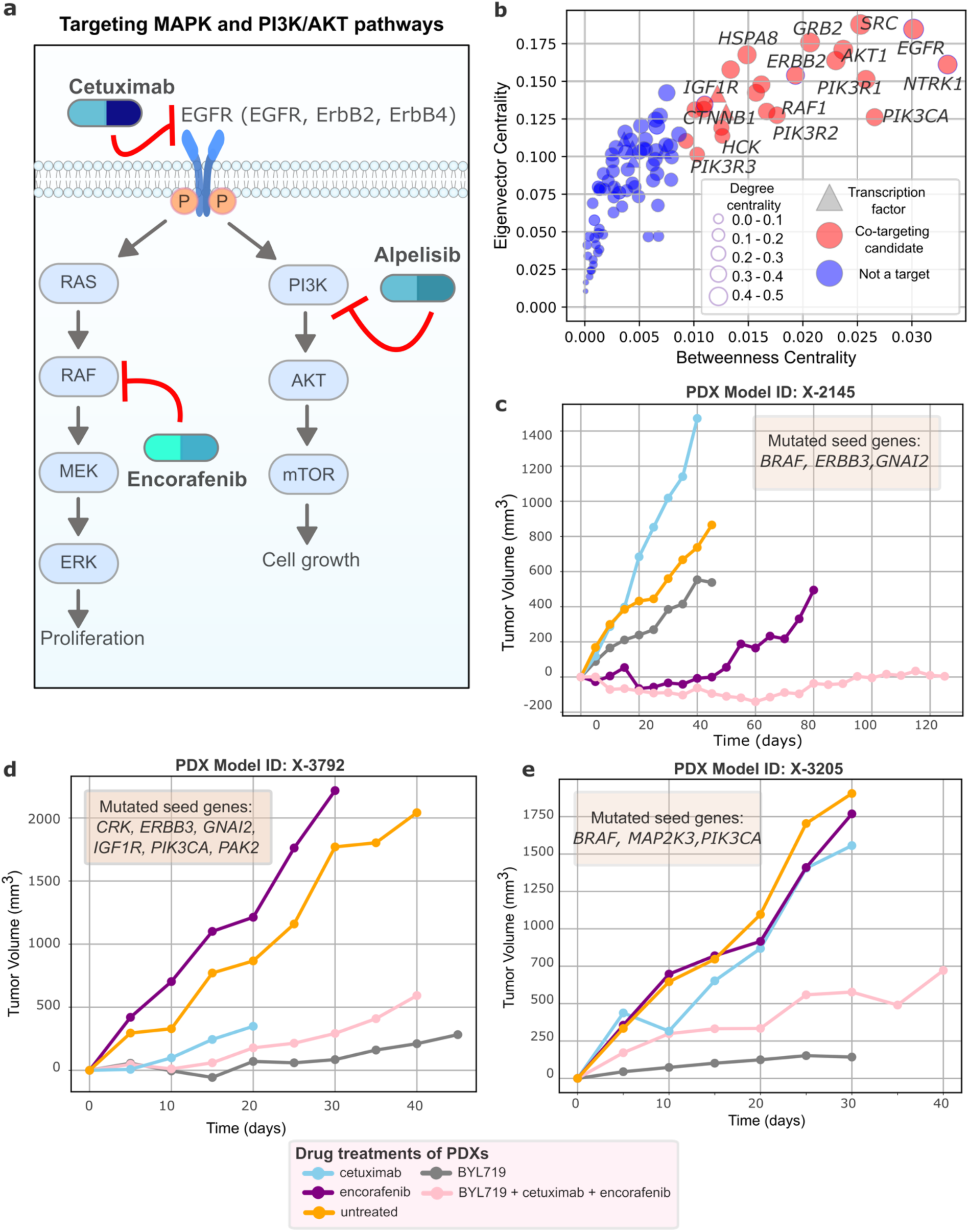
Co-targeting candidates in *BRAF|PIK3CA* subnetwork. **(a)** A simple representation of the MAPK and PI3K/AKT pathways with the nodes mapped from the subnetwork in Figure 6 to delineate the connection between combination therapy targets in a cell. The combination therapy targeting EGFR, RAF and PI3K with cetuximab+encorafenib+alpelisib is shown. In general, if oncogenic signaling is elevated in these pathways through mutations or overexpression, an upstream RTK (*EGFR, ErbB2, ErbB3, ErbB4*) can be combined with downstream oncogenes RAS, RAF, MEK or ERK and PI3K, AKT from the compensatory pathways MAPK and PI3K/AKT, respectively. **(b)** Scatter plot showing the betweenness centrality (x-axis) and eigenvector centrality (y-axis) of the *BRAF*|*PIK3CA* subnetwork in colorectal cancer. Co-targeting nodes (red) on the upper right quadrant of the scatter plot have both high betweenness and eigenvector centrality. These nodes govern important pathways and serve as vital connectors linking sections of the network. The remaining network nodes (blue dots) are not potential targets. Node sizes correspond to their degree centrality, indicating their level of connectivity within the network. **(c-e)** Tumor growth rates of the colorectal cancer PDXs with mutations in the *BRAF|PIK3CA* subnetwork. The growth curves are without treatment, treated with single drugs BYL719 (PI3K inhibitor), cetuximab (EGFR inhibitor), encorafenib (*BRAF* inhibitor) and the combination treatment BYL719+cetuximab+encorafenib. In the models, combination therapy significantly reduces tumor growth rates. x-axis is the treatment days.

For CRC, the RTKs *CSF1R* (encoding CSF-1), *EGFR, ERBB2*, *FLT4, IGF1R*, *PDGFRA/B* can be targeted in combination with MAPK pathway members *SRC* (encoding Src), *RAF1*, *BRAF*, *MAP2K1* (MEK1), *MAP2K2* (encoding MEK2) and PI3K/AKT members *KDR*, *PIK3CA/B/D/G or PRKDC* (PKCδ). These genes have higher betweenness and edge centrality values (Figure 7b) and can thwart the oncogenic signal flow in CRC tumors that have mutations in *BRAF* or *PIK3CA* and at least one of the seed genes.

In Figure 7c we see that the CRC PDX model X-2145 diminishes after the combination of BYL719+cetuximab+encorafenib targeting *PIK3CA* (or PIK3C/B/D/G), *EGFR* and *BRAF*, respectively. The model has mutations on seed genes *BRAF*, *ERBB3* and *GNAI2*. CRC models X-3792 and X-3205 also respond better to the BYL719+cetuximab+encorafenib treatment compared to single treatments (Figure 7d-e). X-3792 has mutations on seed genes *CRK, ERBB3, GNAI2, IGF1R, PIK3CA and PAK2*. X-3205 has mutations on *BRAF, MAP2K3* and *PIK3CA*. For this model, signaling could be stronger as both co-mutated genes have mutations on X-3205, therefore after a small increase followed by a steady state for 20 days, the tumor exhibits a higher growth state. Figures S10 and S11 show the expression levels of some proteins and pathway scores in these PDXs, respectively. *AKT1*, *ERBB2*, *ERBB3* and *MAP2K2* expression levels are higher and above 0.5. As to the expression scores, Thyroid, Rap1, Oxytocin, Hippo, HIF-1 and Estrogen pathways display prominent activity.

This type of combination therapy falls into targeting parallel pathways.^78^ Both MAPK and PI3K/AKT pathways receive signals through an RTK and transduce it downstream (Figure 7a). Single targeting of these molecules often leads to adaptive resistance mechanisms. *BRAF*^V600E^ mutations, present in a subset of colorectal cancers, activate the MAPK, driving proliferation and survival. However, inhibition of *BRAF* alone can result in compensatory activation of EGFR and PI3K/AKT pathways, diminishing treatment response. Similarly, single drug targeting EGFR or PIK3CA can lead to feedback activation of the MAPK pathway or other compensatory survival signals.

Combination therapy mitigates these escape mechanisms by concurrently blocking multiple critical nodes in these intersecting pathways, thereby providing a more comprehensive suppression of tumor cell proliferation and survival. For instance, *BRAF* inhibitors can be complemented by *EGFR* inhibitors to prevent feedback activation of the MAPK pathway, while PI3K inhibitors can further diminish survival signals emanating from the PI3K/AKT pathway. This multi-targeted approach reduces the likelihood of resistance and can lead to more sustained tumor regression.

Resistance mechanisms to these drugs often involve genetic alterations or epigenetic modifications that reactivate the inhibited pathways or alternative survival routes. Secondary mutations in downstream effectors of e.g., BRAF and PI3K, amplification of alternative growth factor receptors, and activation of parallel signaling cascades are common resistance strategies. By targeting *BRAF, EGFR,* and *PIK3CA* simultaneously, the combination therapy not only disrupts the primary oncogenic drivers but also impedes the cancer cells’ ability to adaptively rewire their signaling networks, resulting in a more robust and durable therapeutic response.

## Discussion

Here, we develop a signaling-based method to discover optimal proteins for the oncologist to co-target with drug combinations. This innovative signaling-based approach harnesses nature, mimicking the cancer’s prolife tactics. A fundamental tenet of cancer’s evading anticancer drugs is circumventing the offending anticancer drugs. Since the drugs commonly block pathways, cancer often maneuvers around the blockade through emerging mutations, and (or) altering expression levels, that allow bypassing it. Our signaling- and mutations-informed method aims to capture the cancer’s ploy.

To accomplish this aim we identified potential targets for combination therapy using a network-based approach. Candidates were selected from pivotal subnetworks nodes, characterized by high betweenness centrality, serving as essential communication hubs. As such, they often serve as homeostatic guardians, making them ideal targets for combination therapies aimed at disrupting oncogenic signaling. By analyzing coexisting mutations from our previous study^32^ and their associated subnetworks, we elucidated the principal communication pathways, employing genes on the shortest paths as seeds. This methodology enabled us to determine key oncogenic signaling nodes and essential connector nodes that reveal potential rewiring routes. Betweenness centrality highlights nodes that serve as “bridges” on the shortest paths, critical in network connectivity. We categorized the 170 protein pairs with coexisting mutations based on the pathway enrichment status of their shortest path proteins: pairs where shortest path proteins are enriched in one pathway, two pathways, or three or more pathways, aiming to identify critical bridging nodes in these subnetworks. We propose that tumors with activating mutations in these subnetwork nodes are harbinger of strong oncogenic signals, thus drug combination targets. Their position in the pathway hierarchy matters.

We cluster proteins with high betweenness centrality into three categories with respect to the signaling pathways they belong to: same or redundant, parallel and compensatory pathways.^33,48^ The oncogenes include RTKs—*EGFR, ERBB2, ERBB3, FGFR2, FGFR3, IGF1R*, and *NTRK1*—and transcription factors—*CTNNB1, ESR1, MDM2, MYC*, and *STAT3*. We applied our results to breast cancer and colorectal cancer subnetworks, derived from *ESR1*|*PIK3CA* and *BRAF*|*PIK3CA* protein pairs, which harbor co-existing mutations specific to these tissues, respectively.^32^ We utilized PDX models with mutations in *PIK3CA* (a gene-pair constituent) and at least one mutation in *BRCA1* or *TP53*, or both (shortest path proteins). Breast cancer PDXs exhibited relatively higher expression levels of *AKT1* and *ERBB2*, classifying them as HER2+ breast cancer subtype. They also showed high estrogen pathway activity. After administering the pan-PI3K inhibitor BYL719 (alpelisib) with the monoclonal antibody targeting *ERBB3* (HER3), we observed a dramatic decrease in tumor size in xenografts X-3077 (growth rate: −5.7 mm³/day) and X-1828 (growth rate: 1 mm³/day), while tumor model X-3078 did not grow for 40 days and then resumed growth at a rate of 5.7 mm³/day. In all three xenografts, the combination therapy of BYL719 and LJM716, inhibiting PI3K and HER3 concurrently, significantly decreased *AKT* levels and halted signaling through RTKs *EGFR*, *ERBB2, ERBB3*, and ERα, as *AKT* modulates estrogen signaling in breast cancer.^79,80^ Akt activates ERα-dependent pathways, which are pivotal in neoplastic transformation.^81–83^ ERα is a transcription factor that induces expression of estrogen-responsive genes. Its subcellular localization and stability are essential.^84,85^ In HER2-overexpressing breast cancer cells, PI3K inhibitors can effectively abrogate *AKT* activation but trigger compensatory activation of the ERK pathway. This activation was linked to HER family receptor engagement, including receptor dimerization, phosphorylation, elevated HER3 expression, and adaptor protein binding to HER2 and HER3.^86–89^ ERK pathway activation was mitigated by MEK inhibitors or anti-HER2 monoclonal antibodies and tyrosine kinase inhibitors. Combining PI3K inhibitors with HER2 or MEK inhibitors diminished proliferation, promoted apoptosis, and enhanced anti-tumor efficacy compared to PI3K inhibitors alone. These findings indicate that PI3K inhibition in HER2-overexpressing breast cancer cells induces ERK pathway dependency, suggesting that combined therapeutic approach with anti-MEK or anti-HER2 agents and PI3K inhibitors improves outcomes.

Similarly, concurrent inhibition of *PIK3CA, EGFR*, and *BRAF* with the combination of BYL719, cetuximab, and encorafenib in colorectal cancer PDX models demonstrated greater efficacy compared to single-agent treatments. The monoclonal antibody cetuximab targets EGFR effectively treating metastatic colorectal cancer. EGFR signaling involves KRAS-mediated activation of *BRAF* and *PIK3CA*-driven *AKT1* phosphorylation. *PIK3CA* mutations independently reduce responsiveness to cetuximab in metastatic colorectal cancer, underscoring concurrent inhibition of PI3K/AKT and Ras pathways for treatment outcomes.^90^ Van Geel et al.’s clinical study aligns with our results, supporting combination of *BRAF* and *EGFR* inhibitors to effectively suppress MAPK signaling in *BRAF*^V600E^ colorectal cancer models. Clinical trials in refractory BRAF^V600^-mutant metastatic CRC tested encorafenib plus cetuximab, with or without alpelisib, to establish safe doses. Encorafenib plus cetuximab showed promising clinical activity, with overall response rates of 19% and 18% and median progression-free survival of 3.7 and 4.2 months in dual- and triple-therapy groups, respectively, highlighting combined therapies in BRAF-mutant metastatic colorectal cancer.^91^

The *ESR1|PIK3CA* subnetwork underscores the potential efficacy of inhibiting the PI3K/AKT pathway by targeting an RTK and a downstream kinase such as *PIK3CA/B/D/G*. Concurrently targeting the compensatory pathways MAPK and PI3K/AKT by inhibiting EGFR and the downstream oncogenes *BRAF* and *PIK3CA* is also effective. These co-targeting genes from different clusters in the dendrogram shown in Figure 2, which have a Hamming distance greater than 0.1, can inform combination therapies. This approach involves configuring an upstream RTK with downstream oncogene(s) and/or transcription factors while considering pathway organization and crosstalk. RTKs complexes, as in the case of inhibiting HER3 in HER2+ breast cancer, can guide alternative combinations, as targeting one heterodimer component can inhibit ligand-induced signaling.

### Targeted therapies and single therapy resistance

Targeted therapies can be dichotomized into two major classes: small-molecule inhibitors and monoclonal antibodies, each possessing a distinct mechanism of action.^92–94^ They target proteins critical for cancer progression, enhancing immune responses, disrupting signaling and inducing cell death.^93,95–97^ Monotherapies often fail due to inherent or acquired resistance.^35,78^ Resistance is largely attributed to the high intratumoral heterogeneity, resulting from diverse clonal populations. In breast and colorectal cancers resistance to monotherapy is common. In breast cancer, *PIK3CA* mutations emerge in 30-40% of cases.^12,98,99^ This high mutation load corrupts multiple pathways.^100^ Alpelisib, a targeted treatment, has shown promise when combined with fulvestrant, a hormone therapy, for advanced breast cancer with *PIK3CA* mutations, highlighting the benefit of targeting multiple pathways simultaneously.^21,101^ In colorectal cancer, resistance to treatments like cetuximab, which targets *EGFR*, is often due to *KRAS* mutations.^102,103^ Constitutive *KRAS* activation makes cancer cells resistant to *EGFR* inhibitors. Combination therapies targeting multiple pathways, such as cetuximab with inhibitors to downstream effectors have shown promise.^58,70,104,105^

One of the earliest and most seminal examples of successful clinical translation of targeted molecular therapeutics was the imatinib, a small molecule tyrosine kinase inhibitor (TKI), for treatment of chronic myeloid leukemia (CML) driven by the constitutively active BCR-ABL1 fusion oncoprotein.^79,80,106,107^ Notably, EGFR TKIs are exemplified in anti-non-small cell lung carcinomas (NSCLC) with activating mutations in the *EGFR* gene or gene rearrangements involving *ALK*, *MET*, *NTRK*, and *ROS1* ^81–83^; *BRAF* inhibitors for *BRAF*-mutant melanoma ^108^ and more recently, *KRAS*^G12C^ inhibitors for NSCLC driven by this historically undruggable oncogenic *KRAS* variant.^109,110^ *EGFR* inhibitors are the preferred treatment for lung adenocarcinomas with *EGFR*-sensitive mutations; however, acquired resistance remains a challenge.^111^ When tumors with the secondary T790M mutation are treated with osimertinib, a third-generation *EGFR* inhibitor, resistance often develops primarily due to the C797S mutation.^111–113^ Other resistance mechanisms include *HER2* and *MET* amplification, mutations in *PIK3A*, *KRAS*, and *BRAF*, and epithelial-mesenchymal transition (EMT).^114–116^ Aberrant signaling pathways in malignant cells can result in emergence of resistant phenotypes, which in turn necessitate personalized combination therapies.^35,78,117^

A recent study revealed that metastatic lung adenocarcinoma patients with co-mutations in *EGFR* and *TP53* are more likely to exhibit mixed responses to EGFR TKIs than those with only an EGFR mutation.^45,118–120^ The combination of whole genome doubling (WGD), which drives oncogenic loss of chromatin segregation ^121^ and *TP53* mutations leads to increased genomic instability and copy number aberrations in TKI resistance genes. WGD enhances drug resistance by increasing the likelihood of undergoing copy number changes.

### Exploiting PPI networks for identifying combination therapy targets

Biological networks, particularly PPI networks, can help in blocking oncogenic signaling in cancer cells.^114–116,120,122,123^ By integrating large-scale omics datasets and computational models, context-specific network alterations that drive cancer can be discovered, offering potential therapeutic targets. One prominent algorithm is PageRank, which can identify subnetworks by scoring nodes (genes, proteins, metabolites) based on their network connectivity and importance. Higher PageRank scores establish central elements, revealing critical subnetworks. This method helps pinpoint key components and interactions within complex biological systems.^45,118–120^ Subnetwork reconstruction and topological analysis reveals critical regulators and functional units.^120^ Integration of omics data enables identification of key network features such as hubs and modules.^120,123^ Centrality measures and community detection algorithms elucidate essential proteins and hierarchical organizations in cancer signaling and facilitate discovery of driver mutations and drug targets by mapping context-dependent relationships between oncogenes and their effectors.^122,123^ The ICGC/TCGA Pan-Cancer Analysis of Whole Genomes (PCAWG) Consortium employed network and pathway analyses to scrutinize 2583 whole cancer genomes across 27 tumor types, identifying 93 genes with non-coding mutations clustering into modules of interacting proteins and impacting pathways such as chromatin remodeling, cellular proliferation, and developmental processes.^124^ These analyses can uncover novel cancer genes and genetic alterations, providing profound insights into cancer susceptibilities for targeting.

The topological characteristics of PPI networks provide vital insights into the key components of these systems.^125–127^ Hubs (nodes with high degree) and bottlenecks (nodes with high betweenness) are promising therapeutic targets due to their roles in maintaining network stability and functions.^128^ Beyond centrality metrics, functional enrichment analysis and experimental validation are crucial for verifying the potential of identified proteins as drug targets. Node annotations, biological pathways, and gene expression data are crucial for target selection. REFLECT, an advanced machine learning tool was developed to investigate the concept that recurrent co-alteration signatures may be targeted with tailored combination drugs to enhance preclinical and clinical outcomes.^34^ The significance of network-based methods in identifying combination therapies is underscored by tools like REFLECT to integrate biological data and generate optimal therapeutic strategies. Leveraging the complexity and connectivity within PPI networks, these approaches can uncover critical nodes and pathways, toward highly targeted combination treatments.^129^ Through data mining and network analysis, Hany et al., identified synergistic multidrug combinations, including optimized 3- and 4-drug regimens tailored for ERα+/HER2-/PI3Kα-mutant breast cancer subtypes.^130^ These combinations target ERα along with PI3Kα, p21, and PARP1, demonstrating efficacy in tamoxifen-resistant models and patient-derived organoids, suggesting potential for overcoming current treatment limitations. The Pathway Ensemble Tool (PET) identifies pathways and gene combinations associated with poor prognosis, offering potential targets for combination therapies.^131^

## Conclusions

The mounting number of new targeted therapies has exponentially expanded the therapeutic combinations search space calling for an effective solution to this challenge.^35,78,132^ Selecting optimal small molecule combinations among this vast array renders comprehensive clinical testing impractical. Combinatorial kinase inhibitors aim to disrupt specific resistance signal transduction cascades, inhibiting multiple targets simultaneously.^58,133^ Combination strategies entail blocking of upstream and downstream signaling pathways, exploiting synthetic lethality, and concerted targeting of multiple pathways.^33,35,134^

Network-based, signaling-primed combination therapies represent a transformative paradigm in precision oncology and personalized medicine. This approach leverages molecular interactions within cellular networks to identify synergistic drug combinations targeting multiple oncogenic drivers simultaneously. It addresses the commonly recognized limitations of monotherapies. It offsets compensatory mechanisms and by temporally rotating the combinations, it lessens drug resistance. High-throughput omics data harnessed by sophisticated computational algorithms, is a recipe to therapeutic regimens to the heterogeneous nature of cancers. Within this framework, signaling-learned network-informed combination therapies establish a first step in designing a combinatorial strategy that precisely targets tumor-specific molecular alterations. This approach also expedites drug discovery through the repurposing of existing compounds and the identification of novel synergies.

Here we launch the first concept-based signaling mechanism-informed strategy, which takes up the question of “which signaling pathway and protein to select to mitigate the patient’s expected drug resistance” ^135^ deescalating the massive number of possibilities facing the physician and fitting the solution to the patient status. Applications of our strategy is validated by existing patient-based xenograft models. However, translating these into clinical practice necessitates further rigorous validation in preclinical models and meticulously conducted clinical trials.

## STAR Methods

### KEY RESOURCES TABLE

| Data Source | Data Type |
| --- | --- |
| The Cancer Genome<sup>136</sup><br>Atlas | Tumor sample mutation |
| AACR Project<br>GENIE<sup>137</sup> | Tumor sample mutation |
| KEGG 2019<sup>44</sup> | Signaling pathways |
| MSigDB<sup>56</sup> | Gene ontology |
| HIPPIE PPI Network<sup>41</sup> | Protein-protein<br>interaction network |
| PDX tumor growth<br>and drug response<br>data<sup>46</sup> | Patient derived<br>xenografts tumor<br>growth, mutation and<br>gene expression data |
| OncoKB<sup>138</sup> | The list of oncogenes<br>and tumor suppressor<br>genes |
| CMAP<sup>47</sup> | Drugs and target<br>proteins |
| TRRUST<sup>139</sup> | The list of transcription<br>factors |

| **Software and algorithms** | **Website link** |
| --- | --- |
| Python 3 | https://www.python.org |
| MsigDB | https://www.gsea-msigdb.org/gsea/msigdb |
| Cytoscape | https://cytoscape.org |
| EnrichR | https://maayanlab.cloud/Enrichr/ |
| PathLinker | https://apps.cytoscape.org/apps/pathlinker |
| PageRank<br>algorithhm | https://networkx.org/documentation/stable/reference/algorithms/generated/<br>networkx.algorithms.link_analysis.pagerank_alg.pagerank.html |
| NetworkX | https://networkx.org |
| GSEAPy | https://github.com/zqfang/GSEAPy |
| SciPy | https://scipy.org/ |
| NumPy | https://numpy.org/ |
| matplotlib | https://matplotlib.org/ |
| seaborn | https://seaborn.pydata.org/ |
| Scikit-learn | https://scikit-learn.org/stable/ |
| Python scripts | https://github.com/bengiruken/CombinationTherapyTargets |

### Resource availability

#### Lead contact

Further information and requests for resources should be directed to and will be fulfilled by the lead contact, Bengi Ruken Yavuz.

#### Materials availability

This study did not generate new unique reagents.

#### Data and code availability

This paper analyzes existing, publicly available data. These accession numbers for the datasets are listed in the key resources table.

All original code has been deposited on GitHub and is publicly available as of the date of publication https://github.com/bengiruken/CombinationTherapyTargets. The link is provided in the key resources table.

Any additional information required to reanalyze the data reported in this paper is available from the lead contact upon request.

### Method details

#### Data collection and preprocessing

Somatic missense mutation profiles were sourced from The Cancer Genome Atlas^136^ (TCGA) and the AACR’s Project GENIE^137^ (Genomics Evidence Neoplasia Information Exchange). The TCGA mutation annotation file encompasses data from over 10,000 tumors spanning 33 distinct cancer types, consolidated in the merged MC3 file for comprehensive pan-cancer analysis. Variants with insufficient depth coverage or those identified in normal sample panels were scrutinized for potential germline origin and subsequently excluded prior to integration.

The GENIE mutation dataset (Release 6.2-public) includes data from 65,401 patients and 68,897 tumor samples across 648 Oncotree-classified cancer subtypes. Among the GENIE cohort, 2,930 patients possess multiple tumor barcodes. For analytical purposes, primary tumor barcodes were prioritized when available; otherwise, metastatic or unspecified tumor barcodes were used. Specifically, 2,019 patients had sequenced primary tumors, 757 had sequenced metastatic tumors, and 154 had tumors of unspecified classification.

We utilized HIPPIE^41^, the Human Integrated Protein-Protein Interaction Reference, which offers confidence-scored and functionally annotated human protein-protein interactions (PPIs). HIPPIE aggregates interaction data from 10 source databases and 11 additional studies not fully represented in existing databases. The resulting PPI network comprises 19,485 nodes and 783,182 edges. The network was downloaded from https://www.ndexbio.org using UUID: 89dd3925-3718-11e9-9f06-0ac135e8bacf, version 2.3 (29-APR-2022). After eliminating self-interactions, the PPI network consisted of 778,298 nodes.

For signaling pathway analysis, we obtained the KEGG_2019_Human dataset from the libraries deposited at https://maayanlab.cloud/Enrichr/. This dataset includes 308 pathways and 7,802 genes, with 46 pathways specifically classified as signaling pathways.

#### Protein pairs with co-existing mutations

From the TCGA and AACR GENIE pan-cancer datasets, non-synonymous mutations—including missense, nonsense, nonstop, and frameshift mutations that alter a single protein position—were selected for analysis. Cases with unspecified wild-type or mutant residue names were excluded. Following this filtering process, 9,703 tumors from the TCGA cohort and 57,921 tumors from the GENIE cohort were retained, encompassing a total of 1,631,755 point mutations.

To address sample heterogeneity, variants were pre-filtered based on Variant Allele Frequency (VAF), calculated as the ratio of t_alt_count to t_depth values in the Mutation Annotation File (MAF). Mutations with a VAF greater than 0.125 were retained. This approach identified 3,519 hyper-mutated tumors among 62,567 samples, utilizing a Q3+8 x IQR threshold. Subsequent analysis proceeded with 59,048 non-hyper-mutated samples, spanning 619 cancer subtypes and 33 tissue types (including an ‘OTHER’ category).

The total number of mutations with a VAF greater than 0.125 amounted to 395,801, of which 20,157 were detected in three or more non-hyper-mutated tumor samples. Binary combinations were generated from these 20,157 mutations, creating contingency tables for each possible mutation pair (excluding mutations within the same gene). The co-existing mutations were statistically evaluated using Fisher’s Exact Test. Each mutation pair was assessed with a contingency table [[a,b],[c,d]], where ‘a’ represents tumors with both mutations, ‘b’ those with only the first mutation, ‘c’ those with only the second mutation, and ‘d’ the remaining tumors without both mutations (calculated as 59,048 - (a + b + c)). Statistical significance testing included scenarios where ‘a’ was zero, with multiple testing corrections applied via the Benjamini-Hochberg method. Subsequent analyses focused on co-existing mutations that met a significance threshold of q < 0.3 and were observed in at least three tumors. Components of the significant co-existing mutations were classified as known driver mutations (D) or passenger mutations (P) using the Catalog of Validated Oncogenic Mutations from the Cancer Genome Interpreter.

#### Calculation of the shortest paths between protein pairs

We used 3424 different gene double mutations obtained with the above methodology and deposited in our recent study.^32^ These doublets are harbored by 1284 protein pairs. 46 of these doublets are metastatic breast cancer markers and accumulated on 16 protein pairs. We calculated shortest paths by using PathLinker.^140,141^ PathLinker is a graph-theoretic algorithm designed for reconstructing interactions within a specific signaling pathway. It efficiently identifies multiple short paths within a comprehensive protein interaction network. This method can connect any set of sources to any set of targets within an interaction network. We downloaded the PathLinker algorithm from https://github.com/Murali-group/PathLinker.

We used the constituents of 1284 protein pairs as “sources” and “targets” of the HIPPIE PPI network. PathLinker aims to compute a high-quality reconstruction of these pathways. PathLinker is the computes the k-shortest simple paths in the network from any receptor to any transcription factor. PathLinker accomplishes this through Yen’s algorithm with the A* heuristic, enabling highly efficient computation for very large k values (e.g., 10,000) on networks with hundreds of thousands of edges. PathLinker ranks each interaction in the network based on the index of the first path in which it appears.

We gave the first component of each protein pair as source and the second as the target node with the default parameter k=200 to the PathLinker algorithm to compute the *k* shortest simple paths between a source and target node. A simple path is *a path* with no repeated nodes. PathLinker computed 200 simple shortest paths for 1263 protein pairs, for the remaining 21 protein pairs at least one of the pair constituents is not found in the HIPPI PPI network. The lengths of shortest paths -the number of edges traversed from source to target node varies from one to five. The corresponding data is available in Table S1.

#### Pathway enrichment analysis

After computing 200 shortest paths for 1263 protein pairs, we lumped all the nodes on the shortest paths together for each pair. We dubbed such genes as ‘shortest path proteins.’ The number of nodes forming the shortest paths varies from 81 to 166.

Then by using these shortest path proteins, we conducted gene set enrichment analysis by GSEApy (https://github.com/zqfang/GSEApy) to find out prominent pathways through which the signal flows between two nodes. GSEApy represents a sophisticated implementation of Gene Set Enrichment Analysis (GSEA) in Python/Rust, and functions as a wrapper for the Enrichr tool. GSEA is a computational framework that assesses whether a predefined gene set demonstrates statistically significant and concordant alterations between two distinct biological states. We used the parameter gene_sets=’KEGG_2019_Human’. After getting the enrichment results, we only selected the “signaling pathways.”

After obtaining pathway enrichment results from GSEApy, we kept the significant signaling pathways that the shortest path proteins belong to. There are 1248 protein pairs whose shortest paths are enriched in at least one signaling pathway. We also calculated the fraction of shortest path proteins in each signaling pathway among all shortest path nodes. This would give us the signaling pathway(s) that majority of the shortest path proteins belong to. So, for each pair we selected the signaling pathway(s) that contains more than 20% of the nodes in the shortest paths. For the pairs with shortest path proteins passing through more than one signaling pathway, we labelled the pathways as similar if Jaccard Index .0.2; otherwise, we labeled them as different pathways.

We calculated the shortest paths for 1,263 protein pairs containing 3,424 co-occurring mutations using the PathLinker algorithm (Table S1). Subsequent pathway enrichment analysis of the aggregated shortest path proteins revealed 170 protein pairs, each linked to at least one pathway in which the shortest path proteins showed significant enrichment (refer to Table S2). For each of these 170 protein pairs, we curated a seed gene set by selecting the shortest path proteins associated with the enriched pathways. These seed gene sets were then utilized in the PageRank algorithm to facilitate subnetwork reconstruction.

#### Protein pair specific subnetwork construction from shortest path proteins as seeds in enriched pathways

We applied the PageRank algorithm^45^ from the NetworkX implementation (https://networkx.org/documentation/stable/reference/algorithms/generated/networkx.algorithms.link_analysis.pagerank_alg.pagerank.html) as follows: Upon identifying the pathways with which over 20% of the shortest path proteins for each protein pair are linked, we employed these genes and their corresponding protein pair components (excluding those in enriched pathways) as initial seeds for the PageRank algorithm. The initial scores were assigned as 2 for the protein pair components and 1 for the remaining seed genes, with alpha set to 0.90. To select the subnetwork genes, we established the threshold as the minimum (M) of the scores of Gene 1 and Gene 2 (M = minimum[pagerank_score[Gene1],pagerank_score[Gene2]]). For the protein pairs with total number of selected subnetwork genes is less than or equal to 50, we used a less stringent threshold to 0.2*M not to lose interactions in the PPI network. The preprocessed HIPPIE network, devoid of self-edges, was utilized for this process.

The general formula for PageRank algorithm is as follows:

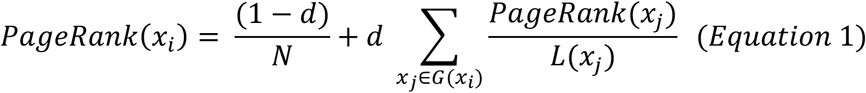

where *x_i_* is a webpage and *x_j_* is a webpage with an outgoing link to *x_i_*.

The PageRank algorithm, a fundamental technique in web-based information retrieval, assesses the relative importance of web pages within a network by evaluating both the quantity and quality of inbound and outbound hyperlinks.^45,142^ It employs a dampening factor, denoted as ‘d’, which ranges between 0 and 1 (empirically calibrated to 0.85 in this study). This factor models the likelihood that a hypothetical user, after navigating through a series of hyperlinks, will randomly access an unlinked page. The algorithm incorporates the cardinality of outbound links from each node, denoted as ‘L’. Essentially, the PageRank algorithm processes an initial set of nodes and produces a ranked list of other nodes in the network, ordered by their relevance to the input set. This is derived from a recursive computation that propagates importance scores through the network’s linkage structure, moderated by the dampening factor and normalized according to each node’s out-degree.

#### Betweenness and eigenvector centrality metrics

Betweenness centrality of a node v is the number of shortest paths passing through that node which can be calculated by the following formula:

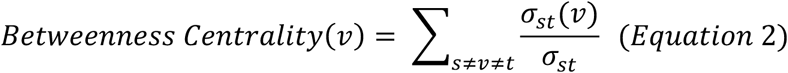

where *σ_st_* is the total number of shortest paths connecting node s to node t and *σ_st_*(*v*) is the total number of shortest paths connecting node *s* to node *t* passing through *v* (https://networkx.org/documentation/stable/reference/algorithms/generated/networkx.algorithms.centrality.betweenness_centrality.html).^143^

To calculate eigenvector centrality, we use the following formula:

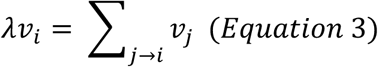

where *λ* is the eigenvalue of maximum modulus that is positive, *v_i_* is the corresponding node (https://networkx.org/documentation/stable/reference/algorithms/generated/networkx.algorithms.centrality.eigenvector_centrality.html).

#### Hierarchical clustering of the connector nodes

We performed hierarchical clustering on a group of genes selected by betweenness centrality metrics. To perform hierarchical clustering, first we divided protein pairs into three groups according to the number of their enriched signaling pathways of the shortest path proteins, the shortest path proteins are enriched in one pathway, two pathways and three or more (multiple) pathways. Then, for the protein pairs we identified the proteins that have betweenness centrality greater than Q3 of their corresponding subnetwork. These are dubbed as the “connector nodes” of the corresponding subnetwork. For each group formed according to pathway enrichment results of the shortest path proteins, we lumped together the connector nodes and then get the intersection of these connector nodes in the three groups. The resulting set of connector nodes in the intersection of the three groups are the ones that are important in cellular network communication.

Then, we created a dendrogram illustrating the relationships between the proteins by evaluating their shared pathways, employing the x-axis to depict the Hamming distance metric that quantify dissimilarity between clusters; smaller distances indicate greater similarity. Greater distances on the dendrogram indicate infrequent co-occurrence of genes within the same signaling pathway, reflecting lower connectivity and functional association. Genes with hamming distance <0.1 can be considered as having similar functions as their corresponding pathways overlap.

Initially, we obtained a catalogue of 46 signaling pathways sourced from KEGG, encompassing a total of 2197 genes distributed among these pathways. Upon analysis of gene-pathway associations, we found genes affiliated with anywhere from one to twelve pathways. Subsequently, we constructed a binary matrix sized at 2197×46, where gene names served as row indices and the 46 pathways as columns. Each cell in the matrix was assigned a value of 1 if the corresponding gene belonged to the pathway specified by the column, and 0 otherwise. Then we calculated the betweenness centrality of the nodes (Equation 2) in each gene-pair specific subnetwork and selected the ones with centrality score >Q3 as connector nodes and clustered them with hierarchical clustering with ‘complete’ parameter.

The Hamming distance, a fundamental metric in information theory and coding theory, serves as a quantitative measure for comparing two binary data strings of equivalent length.^144^ This distance is defined as the cardinality of the set of positions at which the corresponding symbols in two strings are discordant. In the context of our genomic analysis, each gene can be conceptualized as a binary vector of dimension 46 (corresponding to the number of signaling pathways under consideration), where each element is assigned a value of unity if the gene is a constituent of the pathway represented by that dimension, and zero otherwise.

The Hamming distance between two vectors, denoted as H(a,b) where a and b are the vectors in question, can be computed through the application of the XOR operation, symbolized as a ⊕ b. The resultant vector from this operation is then subjected to a summation of its elements, which is equivalent to enumerating the non-zero elements. This process can be formally expressed as:

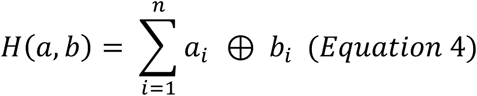

The result of Hamming distance will be the sum of different elements in a series of elements over which distance is compared where n represents the dimensionality of the vectors (in our case, 46).

This metric provides a robust means of quantifying the dissimilarity between genes based on their pathway affiliations, allowing for the construction of a distance matrix that serves as the foundation for subsequent hierarchical clustering analyses.

#### Drug and drug target information

We obtained the data for drugs and their target proteins from the Connectivity map (CMAP).^47^

#### Selection of PDXs for pre-clinical validation

We get the list of PDX models^46^ that have at least one mutated target protein where these proteins also among the shortest path proteins of the corresponding protein pair. We enlisted the seed genes set including source and target nodes; the PDXs should contain at least two drug targets among the seed genes including the source and target nodes. Then, we computed the set of PDX models that have at least two mutations in seed genes for a corresponding given protein pair and also at least one mutation on the protein pair. We also evaluated the volume growth information of untreated tumor, with single treatment and combination therapy.

## Acknowledgments

This project has been funded in whole or in part with federal funds from the National Cancer Institute, National Institutes of Health, under contract HHSN261201500003I. The content of this publication does not necessarily reflect the views or policies of the Department of Health and Human Services, nor does mention of trade names, commercial products, or organizations imply endorsement by the U.S. Government. This Research was supported [in part] by the Intramural Research Program of the NIH, National Cancer Institute, Center for Cancer Research.

## Author contributions

B.R.Y., H.J., and R.N. conceived and designed the study. B.R.Y. did the data curation and visualization, analyzed the results and drafted the manuscript. B.R.Y., H.J., and R.N. validated, reviewed, and edited the manuscript. R.N. supervised the project.

