## Supplemental Information for "Anticancer Target Combinations: Network-Informed Signaling-Based Approach to Discovery"

### **Supplementary Figures**

**Figure S1. Protein pairs where the shortest path proteins are enriched in two or more pathways.** Bubble plots show the signaling pathways where >20% of the pathway genes are enriched in on the y- axis and the corresponding protein pairs harboring co-existing mutations on the x-axis.(**a)** We identified 25 protein pairs where the shortest path proteins enriched in two signaling pathways. Notably, 22 of these protein pairs involve the PI3K/AKT pathway in conjunction with either the MAPK, Ras, Hippo, ErbB, or Chemokine signaling pathways. The pink-highlighted dots represent four metastatic markers: CDH1|PIK3CA, ESR1|PIK3CA, KRAS|PIK3CA, and NUP93|PIK3CA. The shortest path proteins connecting these markers are predominantly enriched in the PI3K/AKT and Ras pathways, with the exception of ESR1|PIK3CA, which shows enrichment in the PI3K/AKT and Chemokine pathways. Protein pairs that traverse three pathways primarily involve the PI3K/AKT, Ras, and MAPK signaling pathways. **(b)** There are 23 protein pairs where the shortest path proteins connecting them traverse three signaling pathways. The node size represents the number of shortest path proteins within the corresponding signaling pathway. The node color indicates the fraction of shortest path proteins in the corresponding pathway among all shortest path proteins connecting the pair. A pink node border denotes that the corresponding protein pair is a breast cancer metastatic marker.

**Figure S2.** There are 91 protein pairs where the >20% of the shortest path proteins pass through only one pathway. The bubble plot below shows the 61 protein pairs where at least one of the components is a drug target in the CMAP data. Dots shows that most of the shortest path proteins connecting the protein pair on the y-axis are enriched in the corresponding pathway on the y-axis.

Each data point reflects the enrichment of shortest path proteins connecting protein pairs along the y-axis within corresponding pathways. Pink data points denote protein pairs linked to breast cancer metastatic markers.

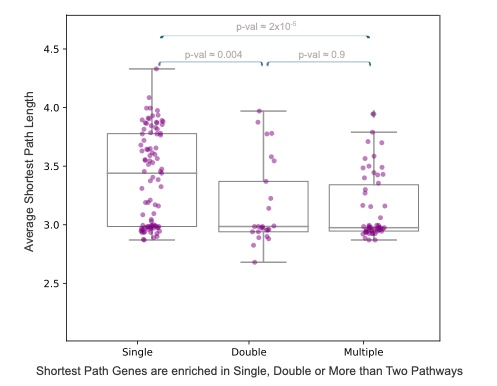

**Figure S3.** For the protein pairs with shortest path proteins enriched in single, double and multiple pathways and the average pathway length for each pair. We compared the average shortest path length distribution of the subnetworks by classifying protein pairs as ‘Single’, ‘Double’ and ‘Multiple’ where the shortest path proteins are enriched in one, two and multiple signaling pathways (Mann-Whitney U test p-values = 0.004 and 2x10-5, respectively). The average shortest path lengths were approximately 3.5 for ’Single’, 3 for ‘Double’, and 3 for ‘Multiple.’ Enhanced crosstalk and coordination between pathways likely facilitate more efficient information flow and signal integration. These findings underscore the adaptive organization of signaling networks, illustrating how interconnected pathways enhance cellular response capabilities.For double and multiple the shortest path length is small because the pathways must be in a cross-talk.

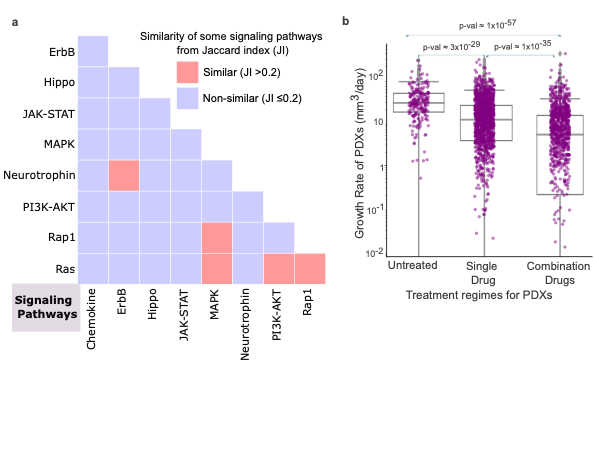

**Figure S4.** Similarity of pathways by Jaccard index similarity metric. Jaccard similarity metric computes a similarity score by $\frac{size(Pathway1\cap Pathway2)}{size(Pathway1\cup Pathway2)}$. If the number of proteins belonging to both pathways is high, the corresponding Jaccard index is high which implies a higher functional similarity between the corresponding pathways.

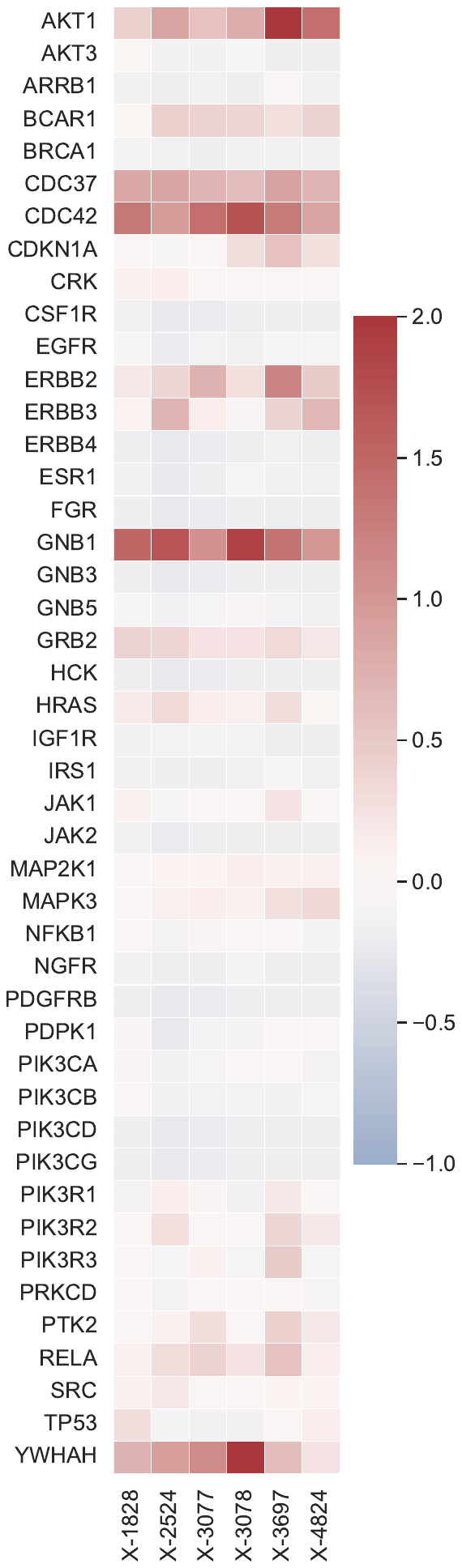

**Figure S5:** Expression values of seed genes and drug targets in BRCA PDX models.

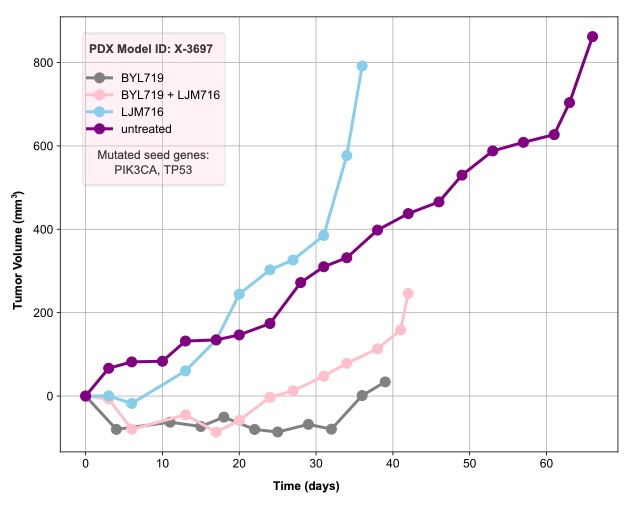

#### **Figure S6.** Growth pattern of PDX model X-3697 exemplifying targeting ESR1-PIK3CA subnetwork in BRCA.

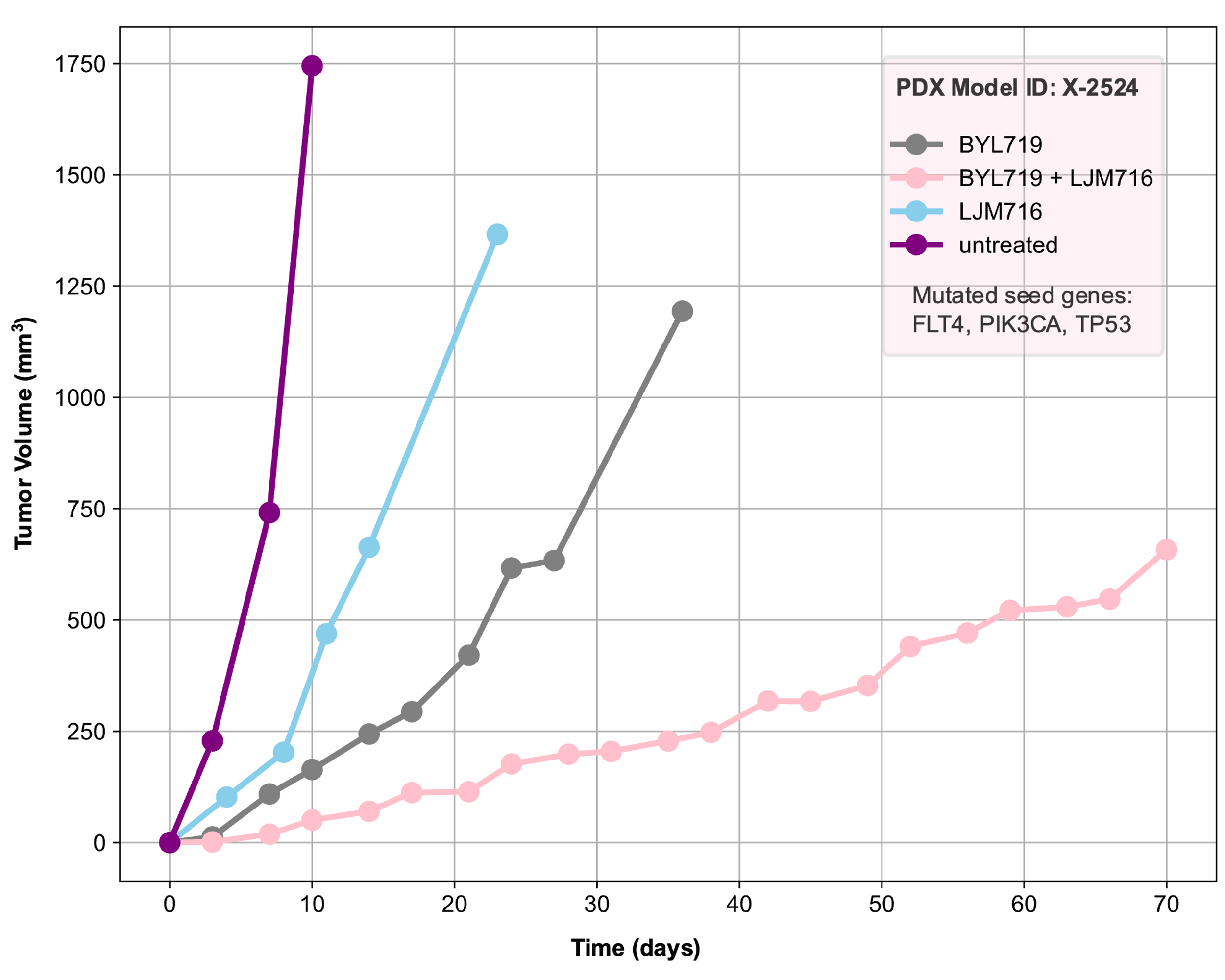

#### **Figure S7.** Growth pattern of PDX model X-2524 exemplifying targeting CDH1-PIK3CA subnetwork in BRCA.

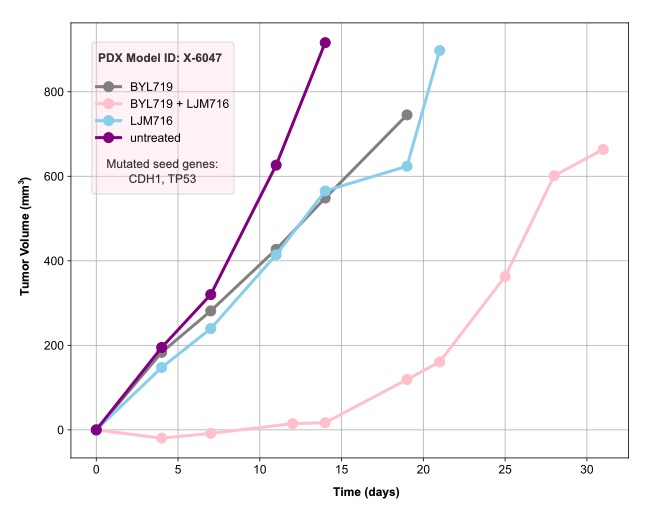

#### **Figure S8.** Growth pattern of PDX model X-6047 exemplifying targeting CDH1-PIK3CA subnetwork in BRCA.

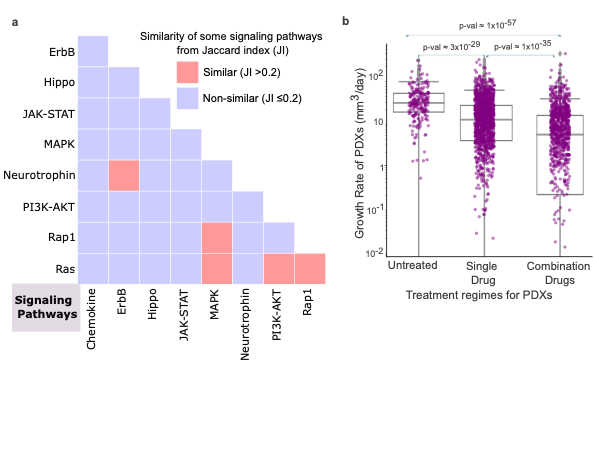

**Figure S9.** Growth rate of the xenografts without treatment, with combination therapies and with single drug treatments.

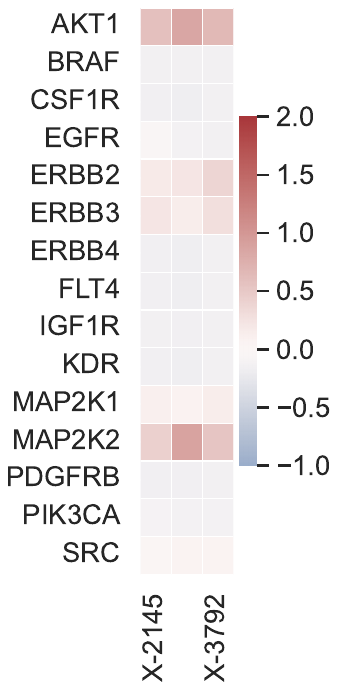

**Figure S10.** Expression values of drug targets in colorectal cancer PDXs having mutations in BRAF-PIK3CA subnetwork.

**Figure S11.** CRC PDXs pathway expression scores for highly changed pathways.
